# A Rapidly Excretable, ROS-Scavenging Ionizable Lipid Decouples mRNA Delivery Potency from Toxicity

**DOI:** 10.64898/2026.04.07.716828

**Authors:** Yeji Lee, Han Jeong, Eunbin Kim, Yuna Hwang, Yongjoo Byeon, Hyunsoo Kang, Min Seok Choi, Eun Hye Jeong, Jeong Hun Kwak, Min-Sung Kang, Ok-Hee Kim, Sairan Eom, Jae Hun Ahn, Yong Jin Lee, Suk Ho Byeon, Say-June Kim, Junwon Lee, Hyukjin Lee

## Abstract

The broader clinical application of mRNA therapeutics remains constrained by dose-limiting toxicities, vector-associated immunogenicity, and prolonged tissue retention of lipid nanoparticles (LNPs) *in vivo*. Here, we report a class of ionizable lipids incorporating a sulfur-bearing hexyl 2-hydroxyethyl sulfide (HHES) motif that decouples mRNA delivery potency from these safety liabilities through dual functionality: the sulfur moiety acts as an intrinsic reactive oxygen species scavenger to suppress oxidative stress, while undergoing oxidative conversion into hydrophilic metabolites to promote rapid systemic clearance. HHES-based LNPs demonstrated a 3.3-fold shorter hepatic half-life and 29-fold lower total hepatic exposure than MC3, while maintaining robust protein expression including functional monoclonal antibody production *in vivo*. Repeated dosing in non-human primates confirmed negligible systemic, hepatic, or hematological toxicity. Leveraging this safety profile, subretinal HHES LNP delivery achieved up to 57% genome editing efficiency in retinal pigment epithelium, suppressing choroidal neovascularization by ∼65% in a wet age-related macular degeneration model without structural damage or microglial activation. This dual-function design provides a generalizable framework for safe, transient, non-accumulative mRNA nanomedicines.

## Introduction

Messenger RNA (mRNA) therapeutics have emerged as a powerful modality for vaccination, protein replacement, cancer immunotherapy, and genome editing. The clinical success of mRNA vaccines and RNA interference therapies has been enabled largely by lipid nanoparticles (LNPs), which protect nucleic acid cargo from degradation and facilitate intracellular delivery [1–3]. Despite these advances, the broader application of mRNA-based therapeutics remains constrained by vector-associated immunogenicity and dose-limiting toxicities, particularly following systemic or repeated administration [4–7].

LNPs are multicomponent systems typically composed of an ionizable lipid, phospholipid, cholesterol, and polyethylene glycol (PEG) lipid [8, 9]. Among these constituents, the ionizable lipid plays a central role in nanoparticle assembly, endosomal escape, and intracellular cargo release [10, 11]. At the same time, accumulating evidence indicates that ionizable lipids are also the primary drivers of LNP-induced innate immune activation and inflammatory responses [12–14]. Commercial and clinically validated ionizable lipids, including DLin-MC3-DMA and SM-102, have been associated with cytokine induction, oxidative stress, and prolonged tissue retention, which can compromise safety margins and limit dosing flexibility [8, 15, 16].

Multiple strategies have been explored to mitigate LNP-associated immunogenicity, including chemical modification of mRNA, incorporation of biodegradable linkages, and rational lipid design to enhance endosomal escape while reducing inflammatory signaling [14, 15]. Recent studies have highlighted that ionizable lipid structure governs not only delivery potency but also intracellular stress responses, including reactive oxygen species (ROS) generation and membrane destabilization [17–19]. However, a molecular design strategy that simultaneously addresses immunogenicity, oxidative stress, and tissue accumulation while maintaining delivery potency has remained elusive [20, 21]. In addition to systemic delivery, local administration of mRNA therapeutics presents distinct safety challenges. In the eye, which is considered an immune-privileged tissue, even transient inflammatory responses or subtle structural damage can result in irreversible functional consequences [22, 23]. While subretinal delivery enables direct access to retinal pigment epithelial cells and photoreceptors, the tolerance of ionizable lipid-based LNPs in this confined and sensitive environment remains incompletely understood [24, 25]. Existing non-viral delivery platforms for ocular genome editing have demonstrated variable efficacy and safety profiles, emphasizing the need for LNP formulations with minimal inflammatory and cytotoxic liability [26, 27].

Here, we report the identification and characterization of a novel ionizable lipid, HHES, that enables high-level protein expression with markedly reduced immunogenicity and toxicity. Through systematic evaluation, we demonstrate that HHES-based LNPs preserve robust mRNA translation while attenuating intracellular ROS generation and inflammatory signaling compared with benchmark ionizable lipids.

We show that HHES LNPs exhibit favorable pharmacokinetic properties, including accelerated systemic clearance and reduced tissue accumulation following intravenous administration in rodents and non-human primates. Importantly, HHES-based formulations display minimal hepatic toxicity, negligible cytokine induction, and excellent tolerability upon repeated dosing. Extending beyond systemic delivery, we demonstrate that HHES LNPs are well tolerated following subretinal administration in mice, with preserved retinal structure, minimal microglial activation, and intact visual function.

Finally, we establish the therapeutic versatility of HHES LNPs by enabling efficient *in vivo* delivery of mRNA-encoded antibodies and genome-editing systems. HHES LNPs support both CRISPR–Cas9-mediated indel formation and adenine base editing in the retina, resulting in effective suppression of pathological choroidal neovascularization. Together, our findings identify HHES as a next-generation ionizable lipid that decouples delivery potency from immunogenicity and toxicity, providing a safer platform for systemic and ocular mRNA-based therapeutics.

## Results

### Design, Synthesis, and Physicochemical Characterization of HHES Lipids

To develop ionizable lipids that combine efficient mRNA delivery with improved biodegradability and intrinsic antioxidant functionality, we designed a new class of sulfur-containing lipids incorporating a hexyl 2-hydroxyethyl sulfide (HHES) motif. The HHES unit was rationally introduced to enable oxidative conversion into more hydrophilic metabolites, thereby facilitating metabolic clearance while maintaining favorable delivery properties (**Fig. 1a**).

**Figure 1.**
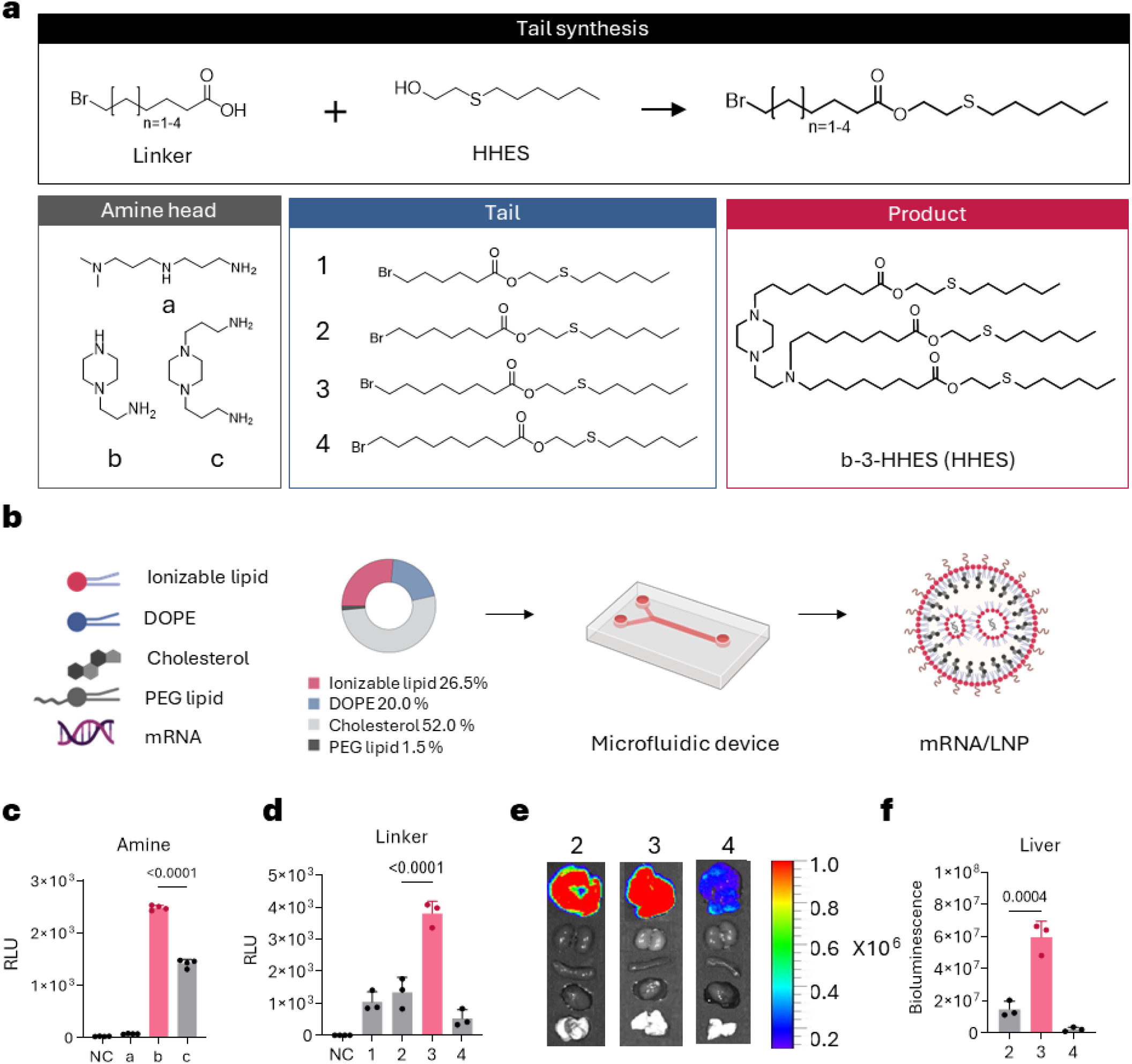
Synthesis and screening of HHES ionizable lipids for efficient mRNA delivery. (a) Schematic overview of the design and synthesis of the HHES lipid library. Ionizable lipids were generated by coupling diverse amine head groups with brominated acid tails of varying lengths. (b) Illustration of LNP formulation using a microfluidic mixing system and the representative lipid composition (ionizable lipid:DOPE:cholesterol:C16-PEG ceramide = 26.5:20:52:1.5). (c) *In vitro* screening of the HHES lipid library with varying amine head groups. LNPs encapsulating fLuc mRNA (20 ng per well) were applied to HeLa cells seeded in white 96-well plates, and luminescence was quantified after 16 h incubation to evaluate the effect of amine head structure on mRNA delivery efficiency (n = 4 per group; one-way ANOVA with post-hoc multiple comparisons: ****p < 0.0001, ***p < 0.001, **p < 0.01, *p < 0.05). (d) Optimization of alkyl linker length within b-series HHES lipids. LNPs encapsulating fLuc mRNA (20 ng per well) were applied to HeLa cells seeded in white 96-well plates, and luminescence was quantified after 16 h incubation (n = 3 per group; one-way ANOVA with post-hoc multiple comparisons: ****p < 0.0001, ***p < 0.001, **p < 0.01, *p < 0.05). (e) *Ex vivo* bioluminescence imaging of major organs 6 h after intravenous injection of fLuc mRNA-loaded LNPs (0.1 mg/kg) in C57BL/6 mice, showing predominant hepatic expression. (f) Quantitative ROI analysis of hepatic bioluminescence across HHES tail-length variants, showing significant differences among groups (n = 3 per group; one-way ANOVA with post-hoc multiple comparisons: ****p < 0.0001, ***p < 0.001, **p < 0.01, *p < 0.05). Data are mean ± SEM.

A combinatorial lipid library was synthesized by coupling diverse amine head groups with brominated acid linkers of varying alkyl chain lengths, allowing systematic modulation of ionization behavior and membrane interaction. All HHES lipids were obtained with high purity and correct molecular composition, as verified by LC–MS and NMR analyses (**Supplementary Figs. 1–9**). Chromatographic and spectroscopic characterization confirmed successful synthesis of the designed structures with minimal impurities, supporting the robustness of the synthetic platform.

The synthesized lipids were formulated into LNPs using a microfluidic mixing system. The optimized HHES LNP formulation consisted of HHES ionizable lipid, DOPE, cholesterol, and C16-PEG ceramide at a molar ratio of 26.5:20:52:1.5. Under these conditions, all HHES LNPs formed uniform nanoparticles with narrow size distribution and high mRNA encapsulation efficiency (>90%) (**Fig. 1b; Supplementary Table 1**). Measurement of ionization behavior using a TNS fluorescence assay showed an apparent pKa of 6.81, indicating suitable protonation characteristics for endosomal escape under acidic conditions (**Supplementary Fig. 10**). Cryogenic transmission electron microscopy revealed spherical particles with multilamellar internal organization, consistent with stable mRNA-loaded LNP formation (**Supplementary Fig. 11**).

### *In Vitro* and *In Vivo* Screening Identifies an Optimized HHES Lipid for Potent mRNA Delivery

We next evaluated the mRNA delivery potency of the HHES lipid library to identify optimal structural features for functional delivery. *In vitro* screening in HeLa cells using firefly luciferase (fLuc) mRNA-loaded LNPs revealed that transfection efficiency was strongly dependent on the amine head group structure (**Fig. 1c).** Among the three head group scaffolds tested, including a linear diamine (N,N-dimethyldipropylenetriamine), a piperazine-based triamine (1-(2-aminoethyl)piperazine), and a bis-piperazine tetraamine (1,4-bis(3-aminopropyl)piperazine), the piperazine-based triamine scaffold (b-series) exhibited the highest luciferase expression, identifying it as the most effective head group for intracellular mRNA delivery.

To further optimize lipid performance, we systematically varied the length of hydrophobic linker within the b-series lipids. The length of alkyl chain significantly influenced delivery potency, with the octanoyl-linked variant (b-3 HHES) producing the highest luciferase expression in HeLa cells (**Fig. 1d**), demonstrating that both head group chemistry and hydrophobic tail architecture critically determine the mRNA delivery efficiency of LNPs.

To validate *in vivo* performance, b-3 HHES LNPs were administered intravenously to C57BL/6N mice at a dose of 0.1 mg/kg fLuc mRNA. Bioluminescence imaging at 6 h post-injection revealed predominant hepatic protein expression (**Fig. 1e**), consistent with the known liver tropism of systemically administered LNPs. Quantitative region-of-interest analysis confirmed significantly higher hepatic luciferase signal for b-3 HHES compared with other tail-length variants (**Fig. 1f**), establishing it as the optimized ionizable lipid for subsequent studies.

### HHES Lipids Exhibit Intrinsic Antioxidant Activity Under Oxidative Stress

Because the HHES ionizable lipid contains a sulfur-bearing moiety designed to modulate ROS, we evaluated its impact on intracellular oxidative stress using a DCFH-DA assay. Cells were first exposed to H₂O₂ (200 μM) for 16 h, followed by treatment with mRNA-loaded LNPs for an additional 16 h. Treatment with SM-102 or MC3 LNPs resulted in a pronounced increase in DCF fluorescence relative to the H₂O₂-only condition (**Fig. 2a**), suggesting that these ionizable lipids further exacerbate oxidative stress, consistent with previously reported membrane-destabilizing effects of conventional ionizable lipids. In contrast, ROS levels maintained comparable to those of the H₂O₂-only control after HHES LNP treatment. At 0.5 μg/mL, SM-102 and MC3 LNPs elevated intracellular ROS levels to approximately 20.10% and 7.18% DCF-positive cells, respectively, whereas HHES LNP-treated cells showed negligible ROS accumulation (∼0.48%) (**Fig. 2b**). At the higher dose of 1.0 μg/mL, SM-102 and MC3 further increased ROS levels to ∼23.16% and ∼18.05%, respectively, while HHES-treated cells remained at ∼1.95%, retaining a markedly lower oxidative burden compared with both benchmark lipids (**Fig. 2c**). Notably, the gap between SM-102 and MC3 narrowed at the higher dose, whereas HHES maintained superior antioxidant activity across both concentrations tested.

**Figure 2.**
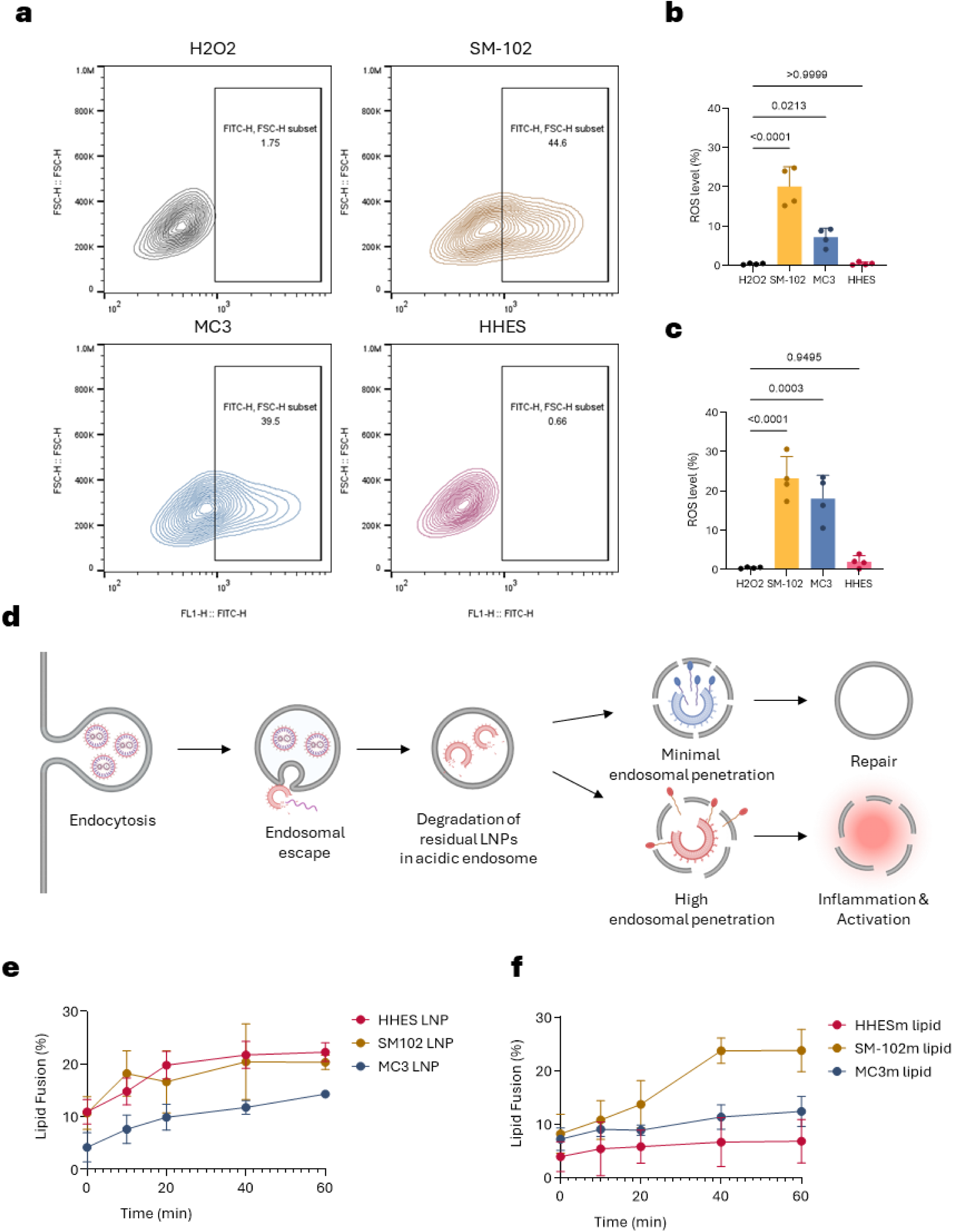
Antioxidant activity and membrane interaction behaviors of HHES LNPs and their ionizable lipid metabolites. (a) Representative flow cytometry plots showing intracellular ROS levels in HeLa cells exposed to H₂O₂ (200 μM) for 16 h, followed by treatment with mRNA-loaded LNPs for an additional 16 h. HHES LNPs markedly reduced DCF fluorescence compared with SM-102, MC3, and H₂O₂-only control groups. (b, c) Quantitative analysis of intracellular ROS levels after treatment with mRNA/LNPs at doses of 0.5 μg/mL (b) and 1.0 μg/mL (c), demonstrating dose-dependent antioxidant effects of HHES lipids (n = 4 per group; one-way ANOVA with post-hoc multiple comparisons: ****p < 0.0001, ***p < 0.001, **p < 0.01, *p < 0.05). (d) Schematic illustration of the proposed intracellular processing and endosomal interaction mechanism of ionizable lipids. Following cellular uptake, ester-containing lipids undergo hydrolysis to generate metabolites, which differentially influence endosomal membrane destabilization. (e) Time-dependent lipid fusion kinetics of HHES, SM-102, and MC3 LNP formulations with endosome-mimicking liposomes measured by a FRET-based assay (n = 4 per group). (f) Fusion efficiency of the corresponding ionizable lipid metabolites (HHESm, SM-102m (heptadecan-9-ol), and MC3m (Dlin-MeOH)), indicating distinct membrane interaction characteristics (n = 4 per group). Data are presented as mean ± SEM.

To confirm the oxidative responsiveness of the HHES motif at the molecular level, we examined chemical changes under oxidizing conditions. FT-IR analysis showed the emergence of a characteristic S=O stretching band after H₂O₂ treatment, consistent with oxidation of the thioether group to sulfoxide and sulfone derivatives, confirming that the HHES sulfur moiety acts as a direct ROS scavenger through oxidative conversion (**Supplementary Fig. 12**). This ROS-suppressing activity was further validated in HepG2 cells, supporting the intrinsic antioxidant capacity of HHES lipids across hepatically relevant cell types (**Supplementary Fig. 13**).

### HHES Lipids and Their Metabolites Modulate Membrane Interaction and Endosomal Processing

The interaction of LNPs with endosomal membranes critically influences the intracellular fate of mRNA cargo and subsequent cellular responses. Following cellular uptake via endocytosis, LNPs are trafficked through early endosomes, where only a limited fraction — estimated at less than 5% of internalized particles — undergoes successful endosomal escape to release mRNA into the cytosol [28, 29]. Because endosomal escape efficiency is inherently low, the majority of internalized LNPs progress to late endosomes, where the increasingly acidic environment promotes degradation of ionizable lipids into smaller metabolites [30, 31]. These degradation products can interact with endosomal membranes, and the extent of membrane perturbation may influence downstream cellular stress responses, including inflammatory signaling [32]. The proposed intracellular processing mechanism is illustrated in **Fig. 2d**.

To characterize membrane interaction behavior, we performed a time-resolved FRET-based lipid fusion assay using endosome-mimicking liposomes. HHES and SM-102 LNPs exhibited comparable lipid fusion kinetics, reaching maximum fusion efficiencies of ∼22.26% and ∼20.33%, respectively, both substantially greater than MC3 LNPs (∼14.31%) (**Fig. 2e**). Because lipid fusion is associated with membrane destabilization events that facilitate endosomal escape, these results are consistent with efficient intracellular mRNA delivery for both HHES and SM-102 formulations.

We next examined membrane interaction of the corresponding ionizable lipid metabolites, HHESm, SM-102m (heptadecan-9-ol), and MC3m (Dlin-MeOH), generated following degradation. In contrast to intact LNPs, metabolite-mediated fusion showed a markedly distinct trend: SM-102m reached ∼23.78%, MC3m ∼12.39%, whereas HHESm exhibited substantially attenuated fusion activity of only ∼6.82% (**Fig. 2f**). The dramatically reduced membrane interaction of HHESm compared with SM-102m and MC3m suggests limited post-degradation membrane perturbation. Given that excessive endosomal membrane disruption has been linked to cellular stress and inflammatory signaling, the attenuated membrane activity of HHES metabolites may contribute to the improved biocompatibility observed with HHES LNPs.

Together, these findings indicate that HHES preserves membrane fusion capacity required for efficient mRNA delivery while minimizing membrane perturbation following lipid degradation, thereby decoupling delivery efficiency from stress-associated downstream effects.

### Rapid Biodistribution and Clearance of HHES Ionizable Lipids

To investigate the *in vivo* biodistribution and clearance behavior of HHES LNPs, we performed quantitative PET/CT imaging following intravenous administration of ⁶⁴Cu-labeled LNPs in C57BL/6N mice. A ⁶⁴Cu-labeled PEG-lipid (⁶⁴Cu–NOTA–DSPE-PEG) was incorporated into the LNP formulation to enable noninvasive tracking of nanoparticle distribution *in vivo* (**Fig. 3a**). Following injection, HHES LNPs rapidly accumulated in the liver, reaching a peak hepatic signal of ∼21.77% ID/g by 4 h post-injection, which progressively declined to ∼14.03% ID/g at 24 h and ∼9.21% ID/g at 48 h, indicating efficient systemic clearance (**Fig. 3b, c**).

**Figure 3.**
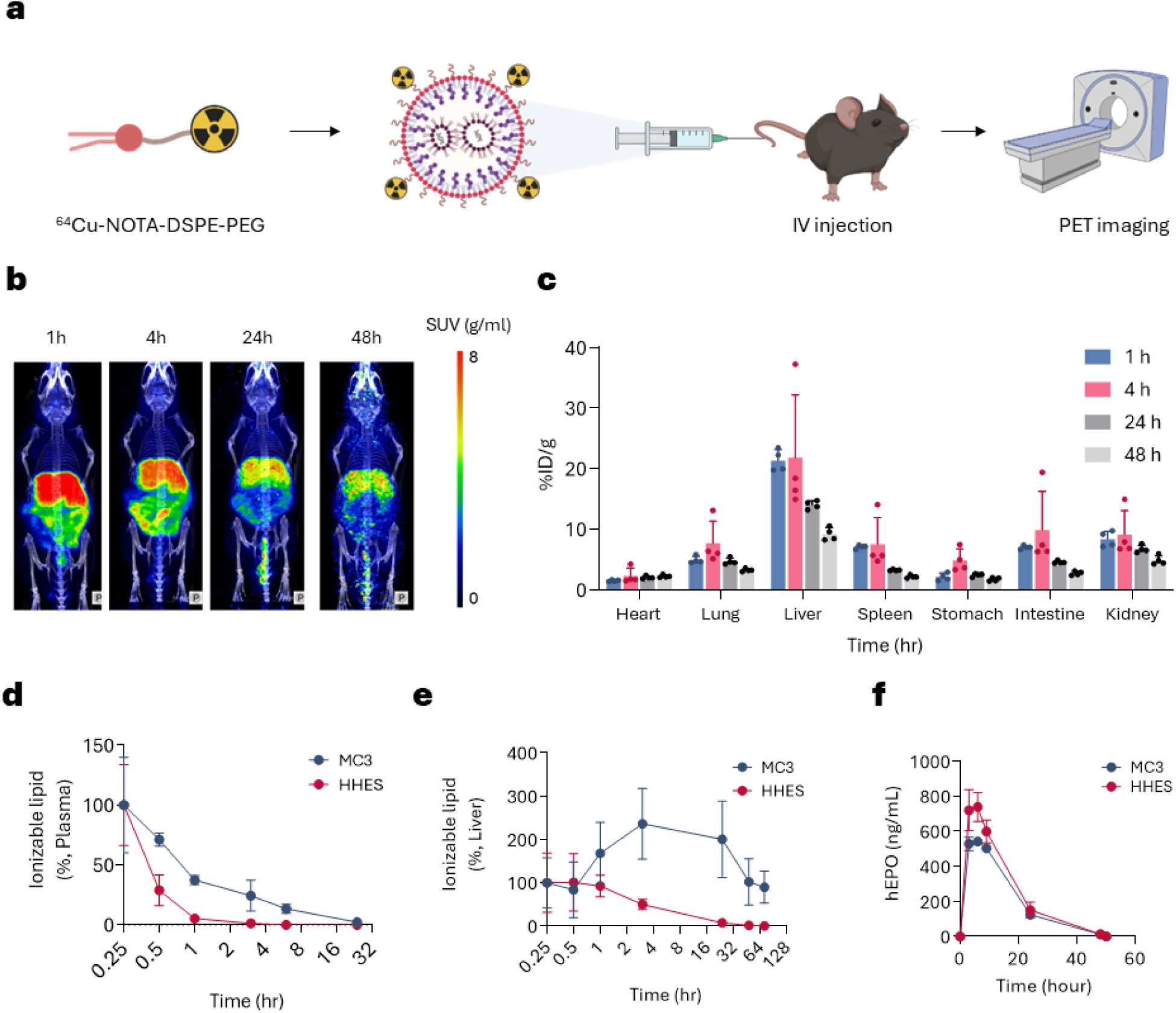
*In vivo* PET/CT imaging, biodistribution, and pharmacokinetics of HHES LNPs. (a) Schematic illustration of the PET/CT imaging workflow for ⁶⁴Cu-labeled LNPs following intravenous injection in C57BL/6N mice. (b) Representative PET/CT images acquired at 1, 4, 24, and 48 h post-injection, showing time-dependent hepatic accumulation and clearance of HHES LNPs. (c) *Ex vivo* biodistribution of ⁶⁴Cu-labeled LNPs in major organs of C57BL/6N mice at 1, 4, 24, and 48 h post-injection, expressed as percentage of injected dose per gram (%ID/g) (n = 4 per group). (d) Plasma pharmacokinetic profiles of HHES and MC3 ionizable lipids in male BALB/c mice (6–8 weeks old) following intravenous administration of mRNA/LNPs (0.5 mg/kg), showing rapid systemic clearance and lower residual plasma levels of HHES compared with MC3 (n = 3 per group). (e) Hepatic pharmacokinetic profiles of HHES and MC3 ionizable lipids quantified in liver tissues by LC–MS/MS in male BALB/c mice (6–8 weeks old) following intravenous administration of mRNA/LNPs (1.0 mg/kg) (n = 5 per group). (f) Serum hEPO expression profiles in BALB/c mice after intravenous administration of hEPO mRNA-loaded HHES or MC3 LNPs (0.1 mg/kg), showing faster onset and higher peak protein expression with HHES compared with MC3 (n = 4 per group). Data are mean ± SEM.

To further characterize the clearance kinetics at the molecular level, we quantified plasma and hepatic ionizable lipid concentrations by LC–MS/MS following intravenous administration of mRNA/LNPs in male BALB/c mice (6–8 weeks old). Plasma pharmacokinetic analysis at 0.5 mg/kg revealed markedly accelerated elimination of HHES compared with MC3: initial plasma concentrations at 0.25 h post-injection were 2,905 ng/mL for HHES versus 17,327 ng/mL for MC3, and HHES levels declined to undetectable levels by 24 h, whereas MC3 remained detectable at 361 ng/mL at the same time point (**Fig. 3d**).

Hepatic pharmacokinetic analysis at 1.0 mg/kg further confirmed the rapid clearance profile of HHES. MC3 exhibited a distinct hepatic accumulation phase, reaching peak concentration (Cmax = 85,774 ng/mL) at 3 h post-injection, with a prolonged half-life of 44.7 h and total hepatic exposure (AUC) of 4,010,006 ng/mL·h. In contrast, HHES reached peak hepatic concentration of only 14,236 ng/mL at 0.5 h post-injection — approximately 6-fold lower than MC3 — and was rapidly eliminated, with a half-life of 13.4 h and AUC of 139,017 ng/mL·h, representing a 3.3-fold shorter half-life and approximately 29-fold lower total hepatic exposure compared with MC3 (**Fig. 3e**).

Together, these results demonstrate that HHES undergoes substantially faster systemic and hepatic clearance than MC3, consistent with its rapid metabolic conversion to hydrophilic species. Notably, unlike conventional ionizable lipids such as MC3 and SM-102, which exhibit transient hepatic accumulation even when biodegradable functionalities are incorporated [15], HHES displayed no evident liver accumulation phase, instead undergoing rapid reduction in hepatic levels following initial uptake, further distinguishing it as a next-generation ionizable lipid with superior clearance properties.

### Rapid Lipid Clearance Does Not Compromise *In Vivo* Functional Delivery

Despite rapid lipid elimination, HHES LNPs maintained efficient functional delivery *in vivo*. Following intravenous administration of hEPO mRNA-loaded LNPs (0.1 mg/kg) in BALB/c mice, HHES induced robust protein expression with rapid onset, achieving peak serum hEPO levels of ∼738.44 ng/mL, modestly exceeding those observed with MC3 (∼542.42 ng/mL) (**Fig. 3f**). These results demonstrate that accelerated clearance does not compromise mRNA delivery efficiency, and may even enhance it.

To further assess delivery at the cellular level, Cre mRNA-loaded LNPs were administered to Ai14 reporter mice, which express tdTomato upon Cre-mediated recombination. HHES induced tdTomato expression in Kupffer cells, hepatocytes, and liver sinusoidal endothelial cells at levels comparable to SM-102, indicating preserved delivery across major liver cell populations (**Supplementary Fig. 14**).

Consistent with these findings, genome-editing studies using Cas9 mRNA/sgRNA-loaded LNPs targeting the *Ttr* locus demonstrated dose-dependent reductions in serum TTR levels comparable to LP-01, an LNP used in Intellia Therapeutics’ clinical gene editing program. Indel analysis further confirmed similar genome-editing efficiencies between HHES and LP-01 (**Supplementary Fig. 15b–d**).

Overall, these results demonstrate that HHES achieves rapid systemic and hepatic clearance while preserving mRNA delivery potency, protein expression, and genome-editing efficacy comparable to established reference LNP platforms.

### *In Vitro* and *In Vivo* Production of Functional Trastuzumab Using HHES LNPs

To evaluate whether the HHES platform supports expression of complex therapeutic proteins, we investigated delivery of mRNAs encoding the heavy and light chains of the monoclonal antibody trastuzumab (**Fig. 4a**). Denaturing agarose gel electrophoresis confirming successful *in vitro* transcription (IVT) of trastuzumab heavy (∼1.8 kb) and light chain (∼1.1 kb) mRNAs, showing single bands corresponding to the expected transcript sizes (Fig. 4b) (**Fig. 4b**).

**Figure 4.**
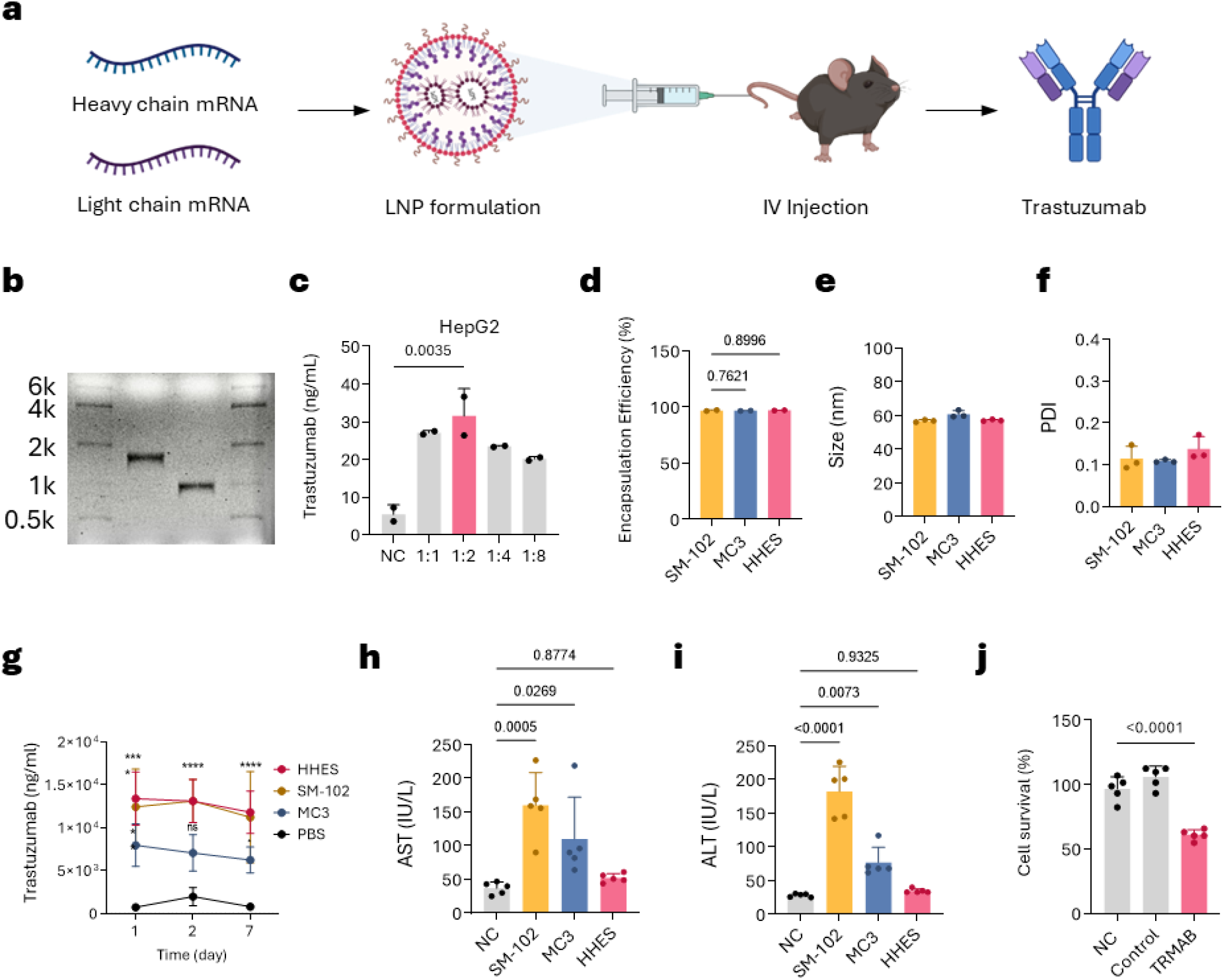
*In vitro* and *in vivo* expression of trastuzumab from mRNA-loaded LNPs. (a) Schematic illustration of the design and *in vivo* delivery strategy of trastuzumab mRNA-loaded LNPs. Separate mRNAs encoding the heavy and light chains of trastuzumab were co-formulated into LNPs and intravenously administered to mice. (b) Denaturing agarose gel electrophoresis confirming successful *in vitro* transcription (IVT) of trastuzumab heavy- and light-chain mRNAs, showing single bands corresponding to the expected transcript sizes. (c) Optimization of the light-to-heavy chain (L:H) mRNA ratio (1:1, 1:2, 1:4, and 1:8) in HepG2 cells (1 μg total mRNA per well), with secreted trastuzumab levels quantified by ELISA (n = 2 per group; one-way ANOVA with post-hoc multiple comparisons: ****p < 0.0001, ***p < 0.001, **p < 0.01, *p < 0.05). (d–f) Physicochemical characterization of trastuzumab mRNA-loaded HHES, SM-102, and MC3 LNPs, including mRNA encapsulation efficiency (d; n = 2), particle size (e; n = 3), and polydispersity index (PDI) (f; n = 3; one-way ANOVA with post-hoc multiple comparisons: ****p < 0.0001, ***p < 0.001, **p < 0.01, *p < 0.05). (g) *In vivo* expression of trastuzumab following intravenous administration of HHES, SM-102, or MC3 LNPs (2 mg/kg total mRNA) in C57BL/6 mice. Plasma antibody concentrations were quantified by ELISA at days 1, 2, and 7 post-injection (n = 4 per group; two-way ANOVA with post-hoc multiple comparisons: ****p < 0.0001, ***p < 0.001, **p < 0.01, *p < 0.05). (h, i) Serum AST (h) and ALT (i) levels measured 24 h post-injection to assess hepatic toxicity (n = 5 per group; one-way ANOVA with post-hoc multiple comparisons: ****p < 0.0001, ***p < 0.001, **p < 0.01, *p < 0.05). (j) ADCC-like assay evaluating the functional activity of mRNA-expressed trastuzumab using HER2-positive BT474 breast cancer cells as target cells and RAW264.7 macrophages as effector cells (n = 5 per group; one-way ANOVA with post-hoc multiple comparisons: ****p < 0.0001, ***p < 0.001, **p < 0.01, *p < 0.05). Data are presented as mean ± SEM.

Optimization of light-to-heavy chain (L:H) mRNA ratios in HepG2 cells identified a 1:2 ratio as optimal, yielding the highest trastuzumab secretion as quantified by ELISA (**Fig. 4c**). Physicochemical characterization demonstrated comparable encapsulation efficiency (∼97%), particle size (∼57–61 nm), and polydispersity index (∼0.11–0.14) among HHES, SM-102, and MC3 LNP formulations, indicating similar nanoparticle quality across lipid systems (**Fig. 4d–f**).

Following intravenous administration (2.0 mg/kg total mRNA) in C57BL/6N mice, HHES LNPs produced sustained plasma trastuzumab levels over 7 days (∼12,400 ng/mL at both day 1 and day 7), whereas SM-102 showed a decline from ∼14,884 to ∼10,036 ng/mL and MC3 exhibited lower overall expression (**Fig. 4g**). Notably, serum AST and ALT levels were markedly lower in the HHES group (AST ∼52 U/L; ALT ∼35 U/L) compared with SM-102 (AST ∼160 U/L; ALT ∼183 U/L) and MC3 (AST ∼110 U/L; ALT ∼77 U/L) (**Fig. 4h, i**), indicating substantially reduced hepatic toxicity associated with the HHES LNPs.

To assess functional antibody activity, an ADCC-like assay was performed using HER2-positive BT474 breast cancer cells and RAW264.7 macrophages as effector cells, which mediate antibody-dependent cytotoxicity via Fcγ receptor engagement. Trastuzumab expressed from HHES-delivered mRNA induced approximately 39% target cell killing compared with untreated controls (**Fig. 4j; Supplementary Fig. 16**), confirming that mRNA-expressed trastuzumab retains functional immune effector activity.

Collectively, these results demonstrate that HHES LNPs enable efficient *in vivo* production of full-length, functional monoclonal antibodies with sustained expression over 7 days, while markedly reducing hepatic toxicity compared with benchmark ionizable lipids, supporting the versatility of the HHES platform for complex protein therapeutics.

### Reduced Hepatic Toxicity and Inflammatory Responses of HHES LNPs in Rats

To evaluate *in vivo* safety following single administration, seven-week-old male Sprague–Dawley rats received a single intravenous injection of mRNA-loaded LNPs at doses of 0.1 or 0.5 mg/kg, and serum biomarkers were analyzed 24 h post-dose. At the higher dose (0.5 mg/kg), SM-102 and MC3 induced marked elevations in AST (∼559 and ∼553 IU/L, respectively) and ALT (∼194 and ∼246 IU/L, respectively), exceeding the normal reference ranges for SD rats (AST: 50–300 IU/L; ALT: 10–80 IU/L). In contrast, HHES maintained AST and ALT within normal ranges (∼198 and ∼42 IU/L, respectively), comparable to control animals (**Fig. 5a, b**). Serum MCP-1 levels were also elevated in SM-102 (∼2,559 pg/mL) and MC3 (∼2,996 pg/mL) groups at 0.5 mg/kg, while HHES-treated animals exhibited markedly lower MCP-1 levels (∼1,478 pg/mL), indicating reduced inflammatory activation following single-dose administration (**Fig. 5c**).

**Figure 5.**
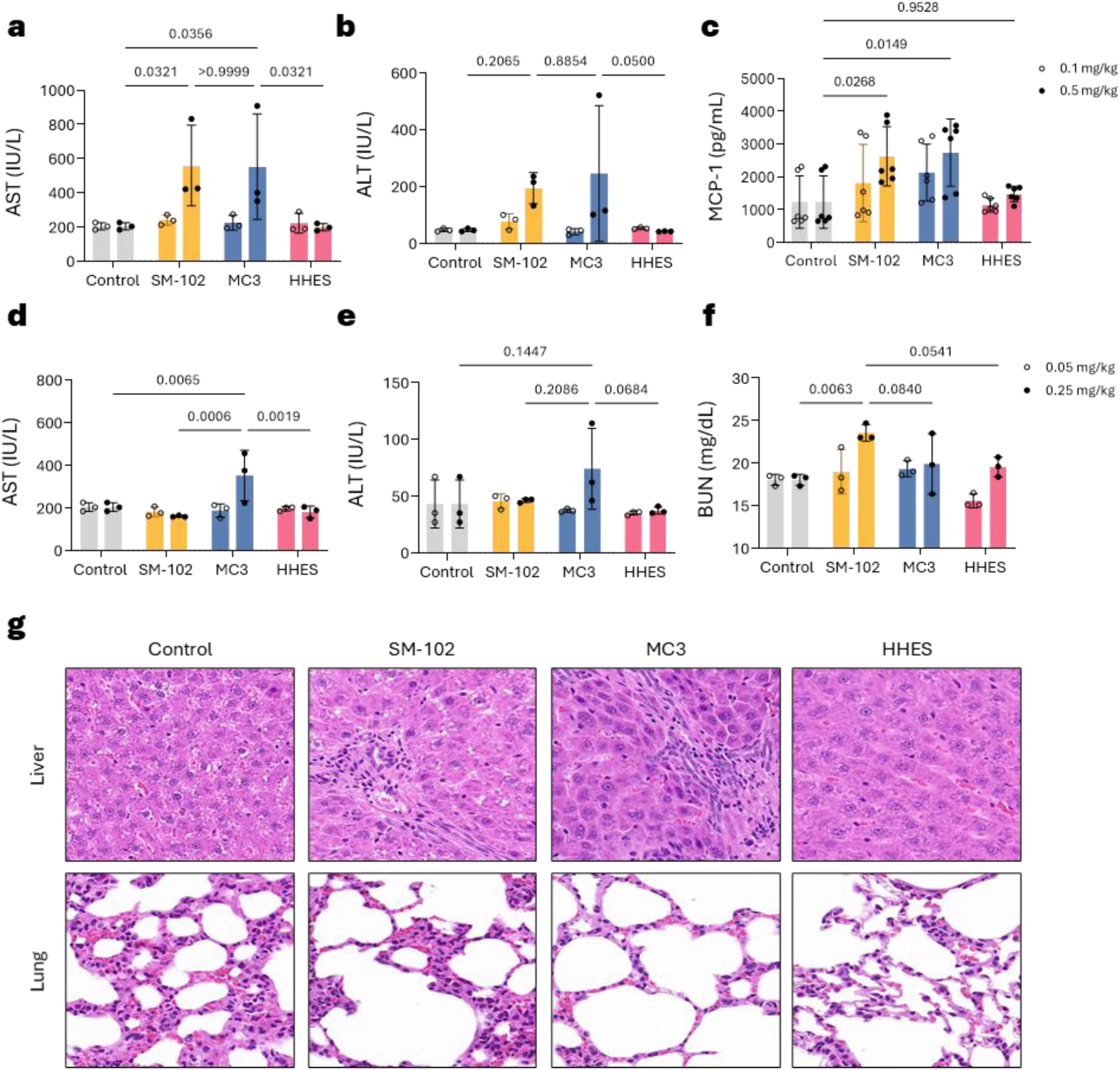
Evaluation of hepatic toxicity and immunogenicity of mRNA-loaded LNPs in rats. (a–c) Serum biochemical and cytokine profiles in seven-week-old male Sprague–Dawley rats 24 h after a single intravenous administration of mRNA/LNPs at doses of 0.1 or 0.5 mg/kg. Serum AST (a), ALT (b), and MCP-1 (c) levels were compared across formulations (n = 3 per group; two-way ANOVA with post-hoc multiple comparisons: ****p < 0.0001, ***p < 0.001, **p < 0.01, *p < 0.05). (d–f) Serum AST (d), ALT (e), and BUN (f) levels following five weekly intravenous injections of mRNA/LNPs at 0.05 or 0.25 mg/kg per dose (n = 3 per group; two-way ANOVA with post-hoc multiple comparisons: ****p < 0.0001, ***p < 0.001, **p < 0.01, *p < 0.05). (g) Representative H&E-stained images of liver and lung tissues collected 24 h after single-dose administration of HHES LNPs at 0.05 mg/kg, showing preserved hepatic architecture and normal alveolar morphology without inflammatory infiltration or necrosis. Scale bars, 100 μm. Data are presented as mean ± SEM.

We next assessed tolerability under repeated dosing conditions. Rats received intravenous injections once weekly for five consecutive weeks at doses of 0.05 or 0.25 mg/kg. At the higher dose (0.25 mg/kg), MC3-treated animals showed a notable increase in AST (∼351 IU/L), approaching the upper limit of the normal reference range, whereas SM-102 (∼160 IU/L) and HHES (∼181 IU/L) remained within normal limits (**Fig. 5d**). ALT levels were similarly well-maintained across all groups, with HHES showing the lowest values (∼37 IU/L) (**Fig. 5e**). BUN levels exhibited a modest increase in the SM-102 group (∼23.6 mg/dL) compared with MC3 (∼19.9 mg/dL) and HHES (∼19.6 mg/dL), suggesting potential renal stress following repeated SM-102 dosing (**Fig. 5f**). Overall, HHES maintained stable biochemical profiles across both doses and repeated administrations.

Histological evaluation was performed 24 h after a single intravenous dose of 0.05 mg/kg mRNA-loaded LNPs. Hematoxylin and eosin staining of liver and lung tissues revealed preserved tissue architecture in HHES-treated animals, with no observable inflammatory infiltration, necrosis, or structural abnormalities (**Fig. 5g**).

Taken together, these findings demonstrate that HHES LNPs induce markedly lower hepatic injury markers and reduced MCP-1–associated inflammatory responses compared with SM-102 and MC3, while maintaining stable biochemical and histological profiles after both single and repeated administration in rats.

### Sustained Protein Expression and Favorable Safety Profile of HHES LNPs in Non-Human Primates

To evaluate translational efficacy and safety, hEPO mRNA-loaded HHES LNPs were administered intravenously to male cynomolgus monkeys (n = 3 per group, Macaca fascicularis, 3–5 years old) at weeks 0, 2, and 4 (**Fig. 6a**). Animals received repeated dosing at doses of 0.05 or 0.25 mg/kg, and serum and hematological parameters were monitored throughout the study.

**Figure 6.**
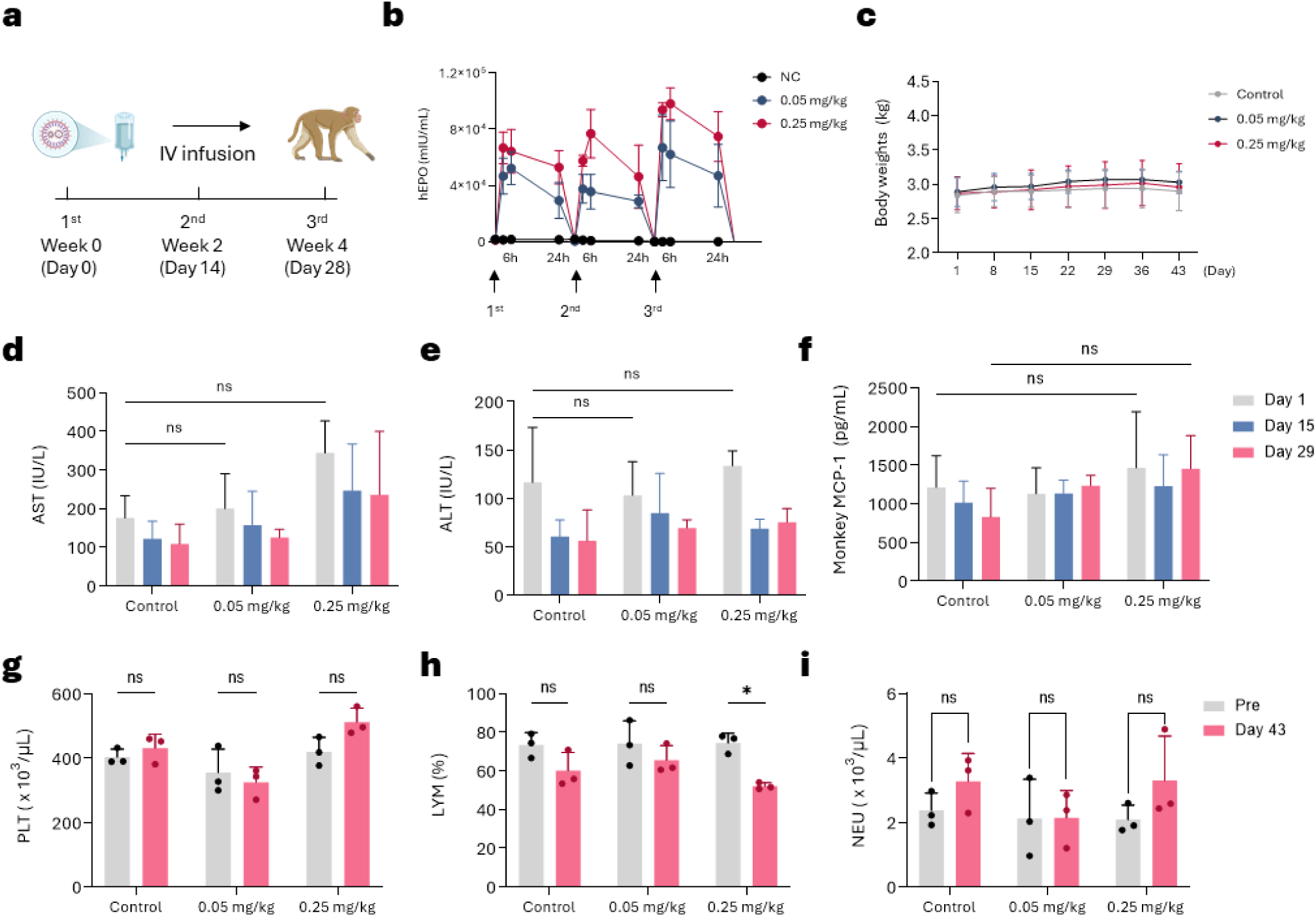
*In vivo* expression and safety evaluation of HHES LNPs in cynomolgus monkeys. (a) Schematic illustration of the dosing regimen for non-human primate (NHP) studies. Male cynomolgus monkeys (n = 3 per group, Macaca fascicularis, 3–5 years old) received intravenous administrations of hEPO mRNA-loaded HHES LNPs (0.05 or 0.25 mg/kg) at weeks 0, 2, and 4. (b) Serum hEPO protein levels measured at 3, 6, and 24 h after each administration, showing transient and reproducible protein expression following repeated HHES LNP dosing (n = 3 per group). (c) Body weight changes monitored throughout the study, showing no significant alterations after repeated administration (n = 3 per group). (d–f) Serum biochemical parameters including AST (d), ALT (e), and MCP-1 (f) measured to assess hepatic function and inflammatory responses (n = 3 per group; two-way ANOVA with post-hoc multiple comparisons: ****p < 0.0001, ***p < 0.001, **p < 0.01, *p < 0.05). (g–i) Hematological parameters including platelet count (g), lymphocyte percentage (h), and neutrophil count (i) evaluated following repeated dosing (n = 3 per group; two-way ANOVA with post-hoc multiple comparisons: ****p < 0.0001, ***p < 0.001, **p < 0.01, *p < 0.05). Data are presented as mean ± SEM.

At both dose levels, HHES LNPs induced robust and reproducible serum hEPO expression following each administration, with peak levels reaching ∼40,000–67,000 mIU/mL at 0.05 mg/kg and ∼67,000–98,000 mIU/mL at 0.25 mg/kg (**Fig. 6b**). EPO expression was consistently observed across all three dosing cycles, demonstrating stable and reproducible *in vivo* delivery performance in non-human primates.

Body weight remained stable throughout the study period at both dose levels, indicating good systemic tolerability (**Fig. 6c**). Serum biochemical analyses revealed no significant changes in AST, ALT, or MCP-1 levels compared with control animals across all dosing cycles (**Fig. 6d–f**), supporting preserved hepatic function and absence of significant inflammatory activation. Renal function markers, including serum creatinine and blood urea nitrogen, similarly remained within normal physiological ranges throughout the study period (**Supplementary Fig. 17**), indicating the absence of treatment-related nephrotoxicity following repeated HHES LNP administration.

Hematological evaluation showed that platelet and neutrophil counts remained stable after repeated administration at both doses, with no significant differences compared with pre-dose values (**Fig. 6g, i**). Lymphocyte percentage at 0.25 mg/kg showed a statistically significant decrease at Day 43 compared with pre-dose values (**Fig. 6h**); however, absolute lymphocyte counts remained within the normal physiological range for cynomolgus monkeys, and this change is likely attributable to the known immunomodulatory effects of supraphysiological EPO levels rather than LNP-associated toxicity.

Consistent with the known pharmacological effects of EPO, modest increases in erythropoietic parameters, including red blood cell count, hemoglobin, hematocrit, and red cell distribution width, were observed following hEPO mRNA administration (**Supplementary Fig. 18**), reflecting expected erythropoietic activity rather than treatment-related toxicity.

Together, these results demonstrate that HHES LNPs enable sustained and reproducible protein expression with minimal systemic, hepatic, or hematological toxicity in non-human primates, supporting their translational potential for repeated systemic mRNA delivery.

Having established that HHES LNPs combine efficient systemic delivery with a favorable safety profile, we next investigated whether these properties extend to local administration in the retina — a tissue where mRNA-based genome editing offers substantial therapeutic potential, yet where even transient inflammatory responses can result in irreversible functional loss.

### HHES LNPs Exhibit Reduced Retinal Toxicity Following Subretinal Injection

To assess ocular tolerability following subretinal injection, wild-type C57BL/6J mice were administered fLuc mRNA-loaded LNPs formulated with HHES, SM-102, or MC3 at 200 or 400 ng/μL, and retinal tissues were analyzed 3 weeks later.

H&E staining showed that retinal laminar architecture was well preserved across all formulations at 200 ng/μL, with no appreciable differences compared with controls. At 400 ng/μL, however, the MC3 group exhibited pronounced structural damage within the region corresponding to the initial subretinal bleb. Specifically, disruption of the RPE monolayer, marked thinning and undulation of the outer nuclear layer (ONL), and undulation of the inner nuclear layer (INL) were observed, accompanied by an overall reduction in retinal thickness at the injection site. The ONL, which normally comprises approximately 10–12 rows of photoreceptor nuclei, was reduced to approximately 4–5 rows in affected areas of MC3-treated retinas. In comparison, SM-102- and HHES-treated retinas did not exhibit overt structural disruption at either concentration, retaining intact RPE monolayer and well-preserved ONL and INL organization (**Fig. 7a**). To quantify photoreceptor layer integrity, retinal thickness and ONL thickness were measured. The area of needle penetration through the neural retina was excluded to avoid confounding by mechanical damage, and measurements were taken at a location 300–500 µm from the needle entry site in the direction of the optic nerve head. At 200 ng/μL, retinal and ONL thicknesses were comparable across all groups and controls. At 400 ng/μL, the MC3 group showed significant thinning of both retinal thickness (98.72 ± 2.98 μm vs. 198.37 ± 3.97 μm in controls) and ONL thickness (1.64 ± 0.90 μm vs. 45.58 ± 0.51 μm in controls; p < 0.0001), whereas SM-102- and HHES-treated retinas showed no statistically significant reduction in either parameter at any measured position (**Fig. 7b**).

**Figure 7.**
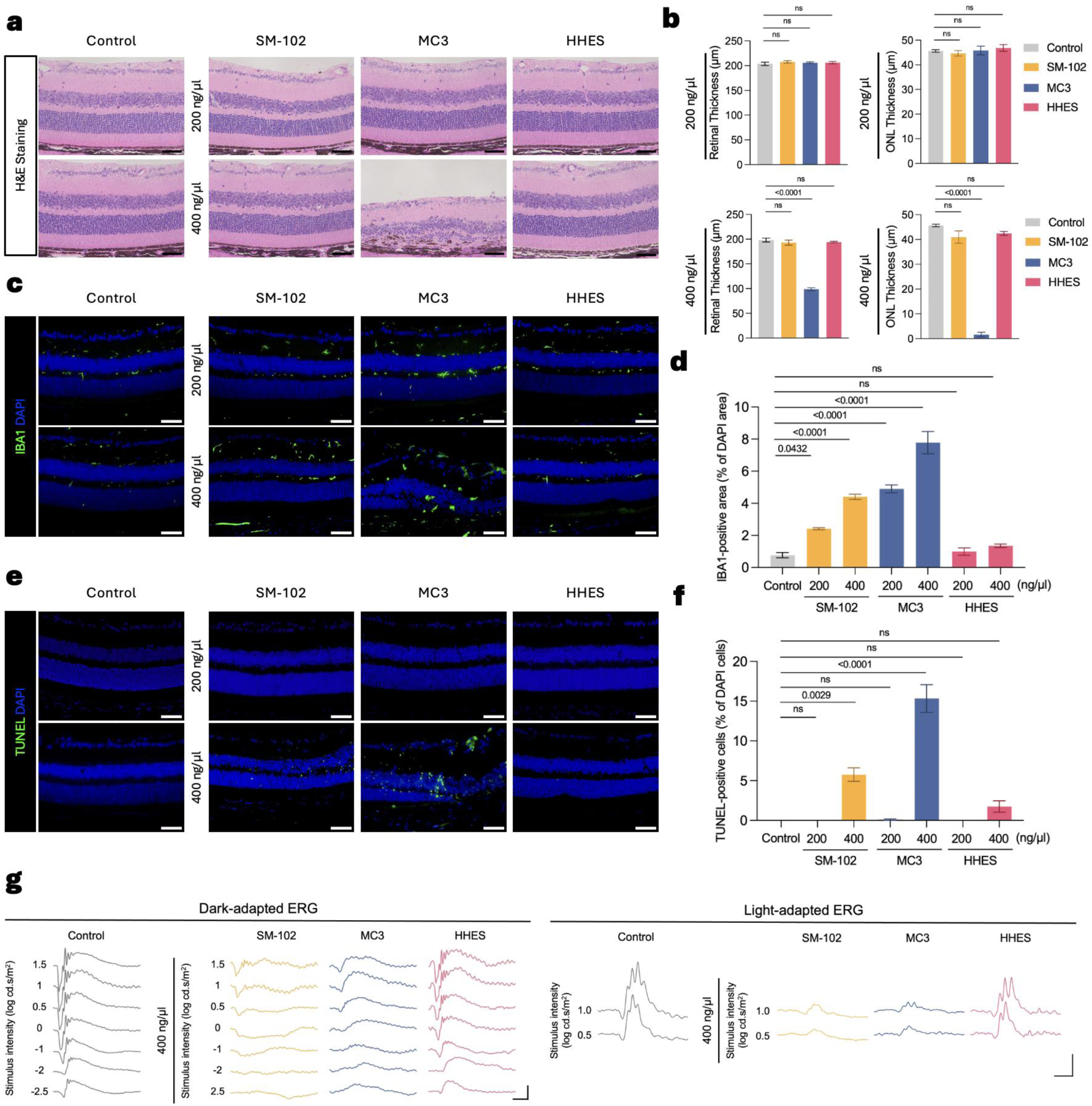
Evaluation of ocular safety following subretinal injection of LNPs in mice. (a) Representative H&E-stained retinal sections collected 3 weeks after subretinal injection of GFP mRNA-loaded LNPs formulated with SM-102, MC3, or HHES at concentrations of 200 or 400 ng/μL in 8-week-old C57BL/6J mice. (b) Quantification of retinal thickness and outer nuclear layer (ONL) thickness at defined locations relative to the injection site. (c) Representative immunofluorescence images of retinal sections stained for IBA1 (microglia/macrophage marker; green) and counterstained with DAPI (blue). (d) Quantification of IBA1-positive area normalized to DAPI-positive area. (e) Representative TUNEL-stained retinal sections (green) with DAPI counterstain (blue). (f) Percentage of TUNEL-positive cells relative to DAPI-positive nuclei. (g) Representative scotopic (dark-adapted) and photopic (light-adapted) ERG waveforms recorded 3 weeks after subretinal injection of LNPs at 400 ng/μL. Scotopic responses were elicited at flash intensities from −2.5 to 1.5 log cd·s/m²; photopic responses at 0.5 and 1.0 log cd·s/m². Data are presented as mean ± SEM. Exact P values from one-way ANOVA with Bonferroni’s multiple-comparisons test are indicated in the graphs. n = 3 biologically independent eyes per group. Scale bars, 50 μm in (a), (c), and (e).

We next evaluated microglial activation and apoptotic cell death in retinal sections. IBA1 immunofluorescence staining revealed minimal microglial activation in controls (IBA1-positive area: 0.77 ± 0.16% of DAPI-positive area) and in HHES-treated retinas at both 200 ng/μL (0.99 ± 0.23%) and 400 ng/μL (1.35 ± 0.11%), with no significant difference between HHES and controls at either concentration. In contrast, the MC3 group showed significantly elevated IBA1-positive area at both 200 ng/μL (4.90 ± 0.25%; p < 0.0001 vs. control) and 400 ng/μL (7.78 ± 0.70%; p < 0.0001 vs. control), and the SM-102 group also showed a significant increase at both 200 ng/μL (2.41 ± 0.07%; p = 0.0432 vs. control) and 400 ng/μL (4.41 ± 0.16%; p < 0.0001 vs. control) (**Fig. 7c, d**). TUNEL staining revealed negligible apoptotic cells in controls and across all formulations at 200 ng/μL. At 400 ng/μL, the MC3 group exhibited the highest percentage of TUNEL-positive cells (15.33 ± 1.75%; p < 0.0001 vs. control), followed by SM-102 (5.74 ± 0.85%; p = 0.0029 vs. control), whereas HHES-treated retinas showed no significant increase relative to controls (1.74 ± 0.71%) (**Fig. 7e, f**).

Beyond structural and molecular assessments of retinal toxicity, we further evaluated retinal function by full-field electroretinography (ERG). Representative scotopic (dark-adapted) and photopic (light-adapted) ERG waveforms recorded at the higher concentration (400 ng/μL) showed markedly attenuated a-wave and b-wave amplitudes in the SM-102 and MC3 groups relative to controls, whereas HHES-treated eyes retained waveform amplitudes comparable to those of controls (**Fig. 7g**).

Taken together, retinal toxicity following subretinal injection showed a dose-dependent trend across formulations. MC3 LNPs elicited the most pronounced effects, including significant microglial activation, apoptotic cell death, structural disruption of the retinal layers, and attenuation of ERG responses. SM-102 LNPs showed intermediate toxicity, with significant inflammatory and apoptotic markers at 400 ng/μL but without overt structural damage. In contrast, HHES LNPs maintained structural, inflammatory, and functional profiles comparable to those of controls at both concentrations tested, demonstrating favorable retinal tolerability consistent with the reduced hepatotoxicity and inflammatory responses observed after systemic administration.

### HHES LNPs Mediate Efficient mRNA Delivery to the RPE

After demonstrating favorable retinal tolerability of HHES LNPs, we next evaluated delivery efficiency and therapeutic potential following subretinal injection. To assess mRNA delivery to the retina, Ai9 reporter mice were treated with HHES-Cre mRNA–loaded LNPs (400 ng/μL) and tdTomato reporter activation was analyzed. Whole-mount imaging of the RPE/choroid complex revealed strong tdTomato fluorescence across a broad area in HHES-treated eyes, whereas no signal was detected in controls. At higher magnification, tdTomato-positive cells displayed the characteristic hexagonal morphology of the RPE, indicating reporter activation within RPE cells **(Fig. 8a).** In retinal sections, tdTomato fluorescence in HHES-treated eyes was predominantly localized to the RPE layer, with no appreciable signal in the neural retina, whereas control retinas showed no detectable signal (**Fig. 8b**). These findings demonstrate that HHES LNPs achieve efficient and selective mRNA delivery to the RPE following subretinal injection.

**Figure 8.**
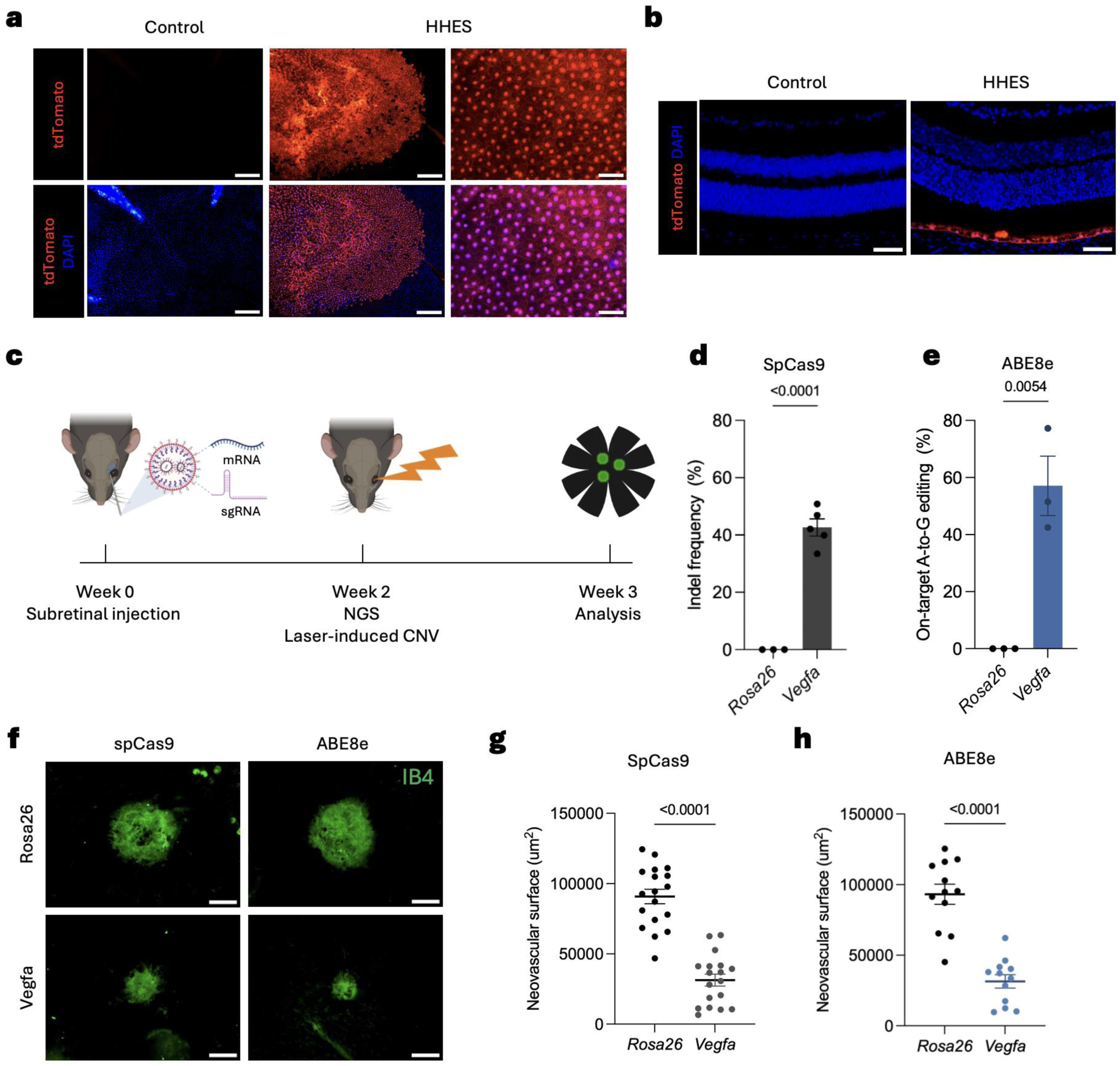
HHES LNP–mediated retinal genome editing suppresses laser-induced choroidal neovascularization following subretinal injection. (a) Whole-mount images of the RPE/choroid complex from Ai9 (LSL-tdTomato) reporter mice showing tdTomato reporter activation after subretinal injection of HHES-Cre mRNA–loaded LNPs (400 ng/μL). Upper panels, tdTomato fluorescence; lower panels, tdTomato merged with DAPI. (b) Retinal cryosections showing tdTomato fluorescence localized to the RPE layer, with no appreciable signal in the neural retina. Counterstained with DAPI. (c) Schematic timeline for subretinal genome editing and evaluation of laser-induced choroidal neovascularization (CNV). HHES LNPs containing SpCas9 or ABE8e mRNA with sgRNA (300 ng/μL total RNA) were injected subretinally at week 0. In a dedicated cohort, RPE cells were isolated at week 2 for targeted deep sequencing. In a separate cohort, laser photocoagulation was performed at week 2 to induce CNV, and lesions were analyzed at week 3. (d) On-target indel frequencies at the *Vegfa* locus following subretinal delivery of HHES LNPs containing SpCas9 mRNA with either *Rosa26* (control) or *Vegfa* sgRNA (n = 3 and 5 eyes, respectively). (e) On-target A-to-G base-editing efficiencies at the *Vegfa* splice donor site following subretinal delivery of HHES LNPs containing ABE8e mRNA with either *Rosa26* (control) or *Vegfa* sgRNA (n = 3 eyes per group). (f) Representative isolectin B4 (IB4)-stained choroidal flat mounts showing CNV lesions in *Rosa26* control and *Vegfa*-targeted eyes. Left columns, SpCas9; right columns, ABE8e. (g) Quantification of CNV lesion area following SpCas9-mediated editing (n = 18 lesions from 6 eyes per group). (h) Quantification of CNV lesion area following ABE8e-mediated base editing (n = 12 lesions from 4 eyes per group). Data are presented as mean ± SEM. P values from unpaired two-tailed Student’s t-test are indicated in (d), (e), (g), and (h). Scale bars, 200 μm and 50 μm in (a), 50 μm in (b), and 100 μm in (f).

Because the performance of gene delivery vehicles may vary across species, we further evaluated whether HHES LNPs can transfect human retinal cells. We first used human ESC-derived retinal organoids, which are primarily composed of neural retina tissue with photoreceptors positioned on the outermost surface; pigmented RPE clusters can form separately and are present on the outer periphery of the organoid. In this configuration, applying LNPs via the culture medium exposes the photoreceptor and RPE surface first, thereby recapitulating the geometry of subretinal injection *in vivo.* Following treatment with HHES-GFP mRNA LNPs (10 μg/mL), GFP-positive cells were detected specifically within the pigmented RPE region of the organoids, whereas no GFP expression was observed in the neural retina compartment (**Supplementary Fig. 19a**). This pattern of RPE-selective transfection was consistent with the *in vivo* findings observed after subretinal injection in mice (**Fig. 8a, b**). We also assessed mRNA delivery in human ESC-derived RPE cells cultured as a monolayer, where LNPs applied in the culture medium access the apical surface of the RPE — the side facing the subretinal space *in vivo* — thereby mimicking the subretinal delivery route. Treatment with HHES-GFP mRNA LNPs at 0.25 and 1.0 μg/mL resulted in readily detectable GFP expression, with both fluorescence intensity and the proportion of GFP-positive cells increasing in a dose-dependent manner (**Supplementary Fig. 19b**). Together, these results demonstrate that HHES LNPs mediate efficient mRNA delivery not only to mouse RPE *in vivo* but also to human RPE cells, supporting the translational relevance of the HHES platform for retinal gene therapy targeting RPE-associated diseases.

### HHES LNP–Mediated *Vegfa* Genome Editing Suppresses Laser-Induced CNV

We next investigated whether HHES LNP–mediated delivery could be harnessed for therapeutic genome editing by targeting *Vegfa* in the RPE, a validated therapeutic strategy for neovascular age-related macular degeneration. Two editing strategies were tested: SpCas9 nuclease to induce targeted indel formation, and adenine base editor ABE8e (TadA-8e fused to Cas9 nickase) [33, 34] to achieve precise base conversion without double-strand breaks. To identify guide RNAs for retinal *Vegfa* editing, we used a previously reported SpCas9 sgRNA targeting exon 3 and selected an ABE8e sgRNA targeting the intron 1 splice donor site using SpliceR v1.3.0 (base-editing window, positions 4–8; ABE score, 0.84), with the aim of disrupting canonical splicing (**Supplementary Fig. 20**).

In mouse Neuro-2a cells, the SpCas9 *Vegfa* guide induced robust indel formation (74.97 ± 0.92%), whereas the *Rosa26* control guide showed no detectable editing. The selected ABE8e guide yielded 40.45 ± 0.65% on-target A-to-G conversion at the *Vegfa* splice donor site, whereas no detectable editing was observed with the *Rosa26* control guide. Consistent with these editing outcomes, western blot analysis showed marked reduction of VEGFA protein, with VEGFA/β-actin ratios of 0.11 and 0.40 for the SpCas9 and ABE8e guides, respectively, relative to the *Rosa26* control (set to 1.0) (**Supplementary Fig. 21**).

To determine whether HHES LNPs can mediate therapeutic genome editing *in vivo*, mice were divided into two cohorts. In the first cohort, RPE cells were isolated two weeks after injection for genomic DNA extraction and targeted deep sequencing to quantify on-target editing efficiency. In the second cohort, laser photocoagulation was performed two weeks after injection to induce choroidal neovascularization (CNV), a widely used preclinical model of neovascular age-related macular degeneration (nAMD), and CNV lesion size was quantified one week after laser induction (three weeks after subretinal injection) (**Fig. 8c**).

Targeted deep sequencing of genomic DNA extracted from isolated RPE cells revealed that HHES LNPs encapsulating SpCas9 mRNA and *Vegfa* sgRNA induced 42.62 ± 2.98% indels at the *Vegfa* locus, compared with 0.00 ± 0.00% in the *Rosa26* sgRNA control group (p < 0.0001) (**Fig. 8d**). Similarly, HHES LNPs encapsulating ABE8e mRNA and *Vegfa* sgRNA achieved 57.07 ± 10.39% on-target A-to-G conversion at the *Vegfa* splice donor site, compared with 0.02 ± 0.02% in the *Rosa26* control group (p = 0.0054), a level consistent with background sequencing error (**Fig. 8e**). These editing efficiencies substantially exceed those previously reported with non-viral delivery platforms in the subretinal space, highlighting the potency of mRNA-based HHES LNP delivery for retinal genome editing. These results demonstrate that HHES LNPs support both SpCas9-mediated indel formation and ABE8e-mediated base editing in the RPE following subretinal injection.

CNV lesion size was next evaluated in the separate cohort. IB4-stained choroidal flat mounts showed reduced CNV lesion size in *Vegfa*-targeted eyes compared with *Rosa26* controls in both the SpCas9 and ABE8e groups (**Fig. 8f**). Quantification of neovascular area showed a significant reduction from 90,754 ± 5,124 μm² in *Rosa26* controls to 31,323 ± 4,218 μm² in *Vegfa*-targeted eyes in the SpCas9 experiment (65.5% reduction; p < 0.0001) (Fig. 8g), and from 93,186 ± 7,127 μm² to 31,449 ± 4,704 μm² in the ABE8e experiment (66.3% reduction; p < 0.0001) (**Fig. 8h**).

Together, these results demonstrate that HHES LNPs enable efficient *Vegfa* genome editing in the RPE, resulting in substantial suppression of pathological neovascularization in a preclinical model of nAMD. The consistent therapeutic effects observed with two mechanistically distinct editing modalities — nuclease-mediated indel formation and base editing–mediated splice disruption — underscore the versatility and translational potential of the HHES LNP platform for ocular gene therapy.

## Discussion

The present study introduces HHES as a dual-function ionizable lipid that simultaneously addresses two persistent liabilities of LNP-based mRNA therapeutics: vector-associated oxidative stress and prolonged tissue accumulation. The sulfur-bearing HHES motif confers intrinsic ROS-scavenging activity through direct thioether oxidation, as confirmed by FT-IR spectroscopy, while its hydrophilic metabolites exhibit markedly attenuated endosomal membrane perturbation compared with those of SM-102 and MC3. This mechanistic combination, in which preserved membrane fusion for endosomal escape is coupled with reduced post-degradation membrane disruption, provides a molecular basis for the observed decoupling of mRNA delivery potency from inflammatory signaling, a challenge that has constrained the clinical application of existing ionizable lipid platforms.

A notable feature of HHES pharmacokinetics is the apparent paradox of accelerated hepatic clearance concurrent with enhanced protein expression. We propose that efficient initial hepatic uptake of HHES LNPs enables productive mRNA delivery, after which rapid oxidative metabolism of the lipid limits cumulative tissue burden without compromising translational output. This interpretation is consistent with the 3.3-fold shorter hepatic half-life, 29-fold lower total hepatic exposure, and absence of a hepatic accumulation phase relative to MC3, together with higher peak serum hEPO levels at equivalent mRNA doses. The safety implications of this clearance profile were borne out in both rat repeated-dose studies and cynomolgus monkey studies, where HHES maintained hepatic enzyme levels, inflammatory markers, and hematological parameters within normal ranges across multiple administrations, a tolerability profile not achieved by benchmark ionizable lipids at comparable doses.

The immune-privileged and anatomically confined nature of the subretinal space places exceptional demands on delivery vector safety, as transient inflammatory responses can produce irreversible photoreceptor loss. The superior retinal tolerability of HHES LNPs relative to MC3 and SM-102, as evidenced by preserved retinal architecture, minimal microglial activation, and intact ERG responses, likely reflects the same attenuated metabolite-mediated membrane perturbation observed *in vitro*. Critically, this safety advantage was not achieved at the cost of delivery efficacy: HHES LNPs supported SpCas9-mediated indel formation at 42.62% and ABE8e-mediated base editing at 57.07% in RPE cells following subretinal injection. These efficiencies compare favorably with previously reported non-viral platforms [35–37]. The editing efficiencies achieved here were sufficient to suppress laser-induced CNV in both SpCas9 and ABE8e cohorts, demonstrating clear therapeutic benefit in a clinically relevant model of neovascular AMD. The consistent therapeutic outcomes across two mechanistically distinct editing modalities underscore the versatility of the platform. RPE-selective transfection was further validated in human ESC-derived RPE cells, which closely recapitulate the morphological and functional properties of native human RPE and have been employed in clinical RPE replacement strategies, supporting the translational relevance of the HHES platform for RPE-associated diseases where human cell validation data remain scarce.

The present study focused on subretinal RPE targeting, which is well suited to the treatment of RPE-associated diseases including wet AMD. Extension of HHES LNP delivery to RPE via intravitreal administration, which would circumvent the surgical complexity of subretinal injection, remains an important direction for future investigation. HHES exemplifies a dual-function design that uses oxidation-labile linkages for simultaneous biodegradability and antioxidant activity. This strategy provides a general framework for developing transient, non-accumulative nanomedicines for various systemic and localized mRNA therapies.

## Materials and Methods

### Materials

Bromo acids, including 6-bromohexanoic acid (150452), 7-bromoheptanoic acid (B3671), 8-bromooctanoic acid (257583), and 9-bromononanoic acid (B2323), were purchased from Sigma-Aldrich (St. Louis, MO, USA) and TCI Chemicals (Tokyo, Japan). Amines and alcohol were obtained from Sigma-Aldrich, including N,N-dimethyldipropylenetriamine (550019), 1-(2-aminoethyl)piperazine (A55209), 1,4-bis(3-aminopropyl)piperazine (239488), and hexyl 2-hydroxyethyl sulfide (S362344). Lipids were purchased from Avanti Polar Lipids (Alabaster, AL, USA) and consisted of 1,2-distearoyl-sn-glycero-3-phosphocholine (DSPC), 1,2-dioleoyl-sn-glycero-3-phosphoethanolamine (DOPE), 1,2-dimyristoyl-rac-glycero-3-methoxypolyethylene glycol-2000 (DMG-PEG 2000, 880151P), and C16-PEG 2000 ceramide (880180). SM-102 (heptadecan-9-yl 8-((2-hydroxyethyl)(6-oxo-6-(undecyloxy)hexyl)amino)octanoate, CAS: 2089251-47-6) was obtained from SINOPEG. Cholesterol (C8667) was provided by Sigma-Aldrich. Heptadecan-9-ol (HY-W099630), MC3 (HY-112251) and Dlin-MeOH (HY-142986) were purchased from MedChemExpress, and LP-01 (Item No. 37278) was obtained from Cayman Chemical.

D-luciferin (P1043) and Bright-Glo™ Luciferase Assay System (E2610) were purchased from Promega. RNA encapsulation efficiency was evaluated using the Quant-iT RiboGreen Assay (Life Technologies, USA). Reporter mRNAs, including firefly luciferase (fLuc), NLS-Cre recombinase (Cre), Cas9, and human erythropoietin (hEPO) (L-7602, L-7211, L-7606, L-7209), were obtained from TriLink BioTechnologies (San Diego, CA, USA). ELISA kits were used to quantify plasma hEPO protein (DEP00; R&D Systems, Minneapolis, MN, USA) and MCP-1 (BMS631INST; Thermo Fisher Scientific, Waltham, MA, USA). Additional reagents, including hyaluronidase (H350.6), deoxyribonuclease I (DNase I; 4716728001), and Liberase Research Grade (5401127001), were supplied by Sigma-Aldrich.

### Synthesis of biodegradable ionizable lipid library

Bromic acid linkers were added to 4 mL of dichloromethane (DCM) containing a mixture of hexyl 2-hydroxyethyl sulfide, DIC, and DMAP. The reaction mixture was stirred at room temperature for 16 h. The resulting carbon tails, bearing both ester and bromine functionalities, were isolated by filtration and further purified using column chromatography on a CombiFlash Rf system with a solvent gradient of 0–10% ethyl acetate in hexane. Subsequently, the carbon tails dissolved in 4 mL of acetonitrile were reacted with an amine structure in the presence of N,N-Diisopropylethylamine. After stirring at 80 °C for 72 h, the solvent was removed under ambient conditions. Final purification was achieved by column chromatography on a CombiFlash Rf system using a gradient of 0–10% methanol in DCM.

### Preparation of ionizable lipid nanoparticles (LNPs)

Lipid nanoparticles (LNPs) were formulated using a microfluidic mixing system (Ignite, PNI, Canada) by combining an ethanol phase containing lipid components with an aqueous RNA phase. Ionizable lipids, cholesterol, helper lipids, and PEG-lipids were dissolved in ethanol. The lipid molar ratios were 26.5:20:52:1.5 for HHES:DOPE:cholesterol:C16-PEG ceramide, 45:9:44:2 for LP-01 and 50:10:38.5:1.5 for SM-102 or MC3:DSPC:cholesterol:DMG-PEG 2000. RNA was diluted in a buffer mixture composed of PBS and 10.0 mM citrate buffer (2:1, v/v). The weight ratio of ionizable lipids to RNA was set at 10:1 for HHES LNPs, 16:1 for MC3 LNPs, 22:1 for LP-01 and 11.5:1 for SM-102 LNPs. Lipid and RNA phases were combined at a 1:3 volume ratio through microfluidic mixing at a flow rate of 12 mL/min. The resulting LNPs were diluted 40-fold in 1× PBS (SH30028.02, Cytiva) and concentrated using Amicon Ultra-15 centrifugal filter units (UFC9010) by ultrafiltration.

### Characterization of LNPs

The size and zeta potential of LNPs were measured after dilution to an RNA concentration of 0.001 mg/mL in 1× PBS. Dynamic light scattering (DLS) was employed to determine the average particle size, polydispersity index (PDI), and zeta potential. RNA encapsulation efficiency was assessed using the Quant-iT™ RiboGreen® Assay (Life Technologies, USA).

### *In vitro* screening of LNP candidates

HeLa cells (ATCC, Manassas, VA, USA) were seeded in white 96-well plates at a density of 1 × 10⁴ cells per well. LNPs encapsulating fLuc mRNA were formulated using microfluidic technology as previously described and applied to the cells at a dose of 20 ng per well. Following a 24 h incubation, cells were lysed with 80 μL of Bright-Glo reagent (Promega) per well, and luminescence intensity was measured using a GloMax® plate reader (Promega).

### *In vivo* screening of LNP candidates

All animal experiments were conducted in accordance with protocols approved by the Institutional Animal Care and Use Committee (IACUC) of Ewha Womans University. For *in vivo* evaluation, LNPs encapsulating fLuc mRNA were prepared as described above and intravenously administered to mice at a dose of 0.1 mg/kg fLuc mRNA. Six hours post-injection, D-luciferin was administered intraperitoneally at a concentration of 6 mg in 0.2 mL. Following a 20 min incubation, luciferase expression was assessed by whole-body imaging and *ex vivo* imaging of harvested organs using an IVIS imaging system (PerkinElmer).

Seven-week-old BALB/c mice (Orient Bio, Seongnam, Korea) were intravenously administered LNPs encapsulating hEPO mRNA at a dose of 0.1 mg/kg. Blood samples were collected via cheek bleeding at 3, 6, 9, 24, and 48 h post-injection, and serum was separated by centrifugation. Serum hEPO protein levels were quantified using an hEPO ELISA kit (R&D Systems) following a 1:1000 dilution of the samples.

### Ionizable Lipid Pharmacokinetics (PK) in Plasma

The pharmacokinetics of HHES and MC3 ionizable lipids were evaluated in male BALB/c mice (6–8 weeks old). For plasma PK, mice were intravenously injected with mRNA/LNPs at a dose of 0.5 mg/kg via the retro-orbital vein. Blood samples were collected at 0.25, 0.5, 1, 3, 6, 24, 48, 72, and 120 h post-injection through the jugular vein using heparinized syringes. Plasma was separated by centrifugation at 2,000 × g for 10 min at 4°C. For hepatic PK, mice were administered mRNA/LNPs at a dose of 1 mg/kg via the retro-orbital vein, and liver tissues were harvested at 0.25, 0.5, 1, 3, 24, 48, and 72 h post-injection. Ionizable lipid concentrations in plasma and liver tissues were quantified using LC–MS/MS. Calibration curves for HHES and MC3 were established using known concentrations to accurately determine lipid levels.

### *In Vivo* PET/CT Imaging and Biodistribution of ⁶⁴Cu-LNPs

For PET/CT imaging, C57BL/6N mice were intravenously injected with approximately 300 μCi of ⁶⁴Cu-labeled LNPs. PET scans were performed at 1, 4, 24, and 48 h post-injection using a small-animal PET/CT system, and reconstructed images were analyzed to evaluate the *in vivo* distribution of LNPs.

For biodistribution studies, mice received ∼30 μCi of ⁶⁴Cu-LNPs via intravenous injection. At 1, 4, 24, and 48 h post-injection, major organs including the heart, lung, liver, spleen, stomach, intestine, and kidney were harvested and weighed. Radioactivity was measured using a gamma counter, and the data were expressed as the percentage of injected dose per gram of tissue (%ID/g).

The LNP formulation consisted of HHES:DOPE:cholesterol:DSPE-PEG-NOTA:C16-PEG ceramide at a molar ratio of 26.5:20:52:0.5:1.

### Assessment of Liver Toxicity and Immunogenicity in rats Liver toxicity

Acute liver toxicity was evaluated in Seven-week-old Sprague-Dawley (SD) rats (n=3) after intravenous administration of mRNA/LNPs at doses of 0.1 or 0.5 mg/kg. Blood samples were collected 24 h post-injection from the jugular vein using heparinized syringes under isoflurane anesthesia. Serum was obtained by centrifugation at 2,000 × g for 10 min at 4 °C. Liver enzyme activities, including aspartate aminotransferase (AST) and alanine aminotransferase (ALT), were quantified using a clinical chemistry analyzer according to the manufacturer’s instructions. Measured values were compared with normal reference ranges for SD rats (AST: 50–300 U/L; ALT: 10–80 U/L).

### Immunogenicity

Systemic immune responses were assessed by measuring serum cytokine levels of MCP-1 using ELISA kits (Thermo Fisher Scientific and R&D Systems). Blood samples were collected 6 h post-injection following intravenous administration of mRNA/LNPs at doses of 0.1–0.5 mg/kg, and cytokine concentrations were expressed as pg/mL.

### Histopathology

For histological analysis, liver and lung tissues were harvested 24 h post-injection of mRNA/LNPs. Samples were fixed in 10% neutral buffered formalin, embedded in paraffin, sectioned at 4–5 μm, and stained with hematoxylin and eosin (H&E). Tissue morphology was examined under light microscopy to assess pathological changes.

### Non-Human Primate (NHP) Study

Male Cynomolgus monkeys (n=3, Macaca fascicularis, 3–5 years old) were used for *in vivo* evaluation of mRNA/LNPs. Animals were intravenously infused with HHES LNPs encapsulating hEPO mRNA at doses of 0.05 mg/kg and 0.25 mg/kg, with administrations performed at weeks 0, 2, and 4 (total of three injections). Control animals received vehicle only.

### hEPO expression

Blood samples were collected at 3, 6 and 24 h following each administration. Serum hEPO protein levels were quantified using ELISA kits (R&D Systems) according to the manufacturer’s instructions, and results were expressed in mIU/mL.

### Serum biochemistry

Blood was also collected prior to each injection and at 24 h post-injection to assess acute toxicity. Serum was separated by centrifugation at 2,000 × g for 10 min at 4 °C. Biochemical parameters, including aspartate aminotransferase (AST), alanine aminotransferase (ALT), blood urea nitrogen (BUN), and creatinine (CRE), were measured using a clinical chemistry analyzer (Hitachi or Roche Cobas system). Hematological parameters, including platelet count, lymphocyte count, and neutrophil count, were assessed using an automated hematology analyzer.

### Reactive Oxygen Species (ROS) Assay

ROS levels were evaluated in RAW264.7 cells using the DCFH-DA fluorescence assay. Cells were seeded in 24-well plates and incubated for 16–24 h prior to treatment. Oxidative stress was induced by adding hydrogen peroxide (H₂O₂, 200 μM; Sigma-Aldrich, Cat. No. 216763) in culture medium. After an additional 16–24 h incubation, cells were treated with LNPs encapsulating mRNA at doses equivalent to 0.5 or 1 μg mRNA per well. Following a further 16–24 h incubation, cells were incubated with 10 μM DCFH-DA (D6883, Sigma-Aldrich) in PBS for 30 min at 37 °C. Intracellular ROS levels were visualized by fluorescence microscopy, and GFP fluorescence intensity was quantified to assess ROS scavenging activity.

### Lipid Fusion FRET Assay

A fluorescence resonance energy transfer (FRET) assay was performed to evaluate lipid fusion between LNPs and endosome-mimicking liposomes. Liposomes were prepared with DOPS:DOPC:DOPE:NBD-PE:N-Rh-PE at a molar ratio of 25:25:48:1:1 using a microfluidic mixing system (flow rate ratio 3:1; total flow rate 12 mL/min.

For the assay, 50 μL of citrate buffer was dispensed into each well of a black 96-well plate, followed by the addition of 1 μL of liposomes (1 mM). LNPs were added to the sample wells at an mRNA concentration of 20 μg/mL in 100 μL volume. Negative control wells (F_min_) contained PBS and liposomes, while positive control wells (F_max_) contained 2% Triton X-100 and liposomes. Plates were incubated at 37 °C for 5 min, followed by 10 s shaking. Fluorescence was measured using a plate reader at λ_ex = 465 nm and λ_em = 520 nm.

Lipid fusion efficiency was calculated according to the equation:

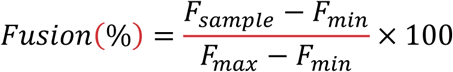

Metabolite assays were performed by preparing metabolite stock solutions (10 mg/mL) in PBS and diluting to the working concentration (100 μg/mL) prior to addition to liposomes.

### IVT mRNA Synthesis for Trastuzumab

DNA templates encoding the heavy and light chains of trastuzumab were designed based on previously reported sequences and synthesized by Integrated DNA Technologies (IDT) [38]. The vector backbones were amplified using a PCR kit (Takara, Japan) to generate templates containing a T7 promoter and poly(A) tail.

*In vitro* transcription (IVT) was carried out using the MEGAscript™ T7 Transcription Kit (Thermo Fisher Scientific, USA) according to the manufacturer’s instructions. The resulting mRNAs were designed to incorporate a Cap1 structure at the 5′ end and a ∼120-nt poly(A) tail. Following transcription, mRNAs were treated with DNase and purified prior to formulation into LNPs.

### *In Vitro* Expression of Trastuzumab

HepG2 cells were seeded in 24-well plates and transfected with LNPs encapsulating trastuzumab heavy and light chain mRNAs (1 μg total mRNA per well). Different light-to-heavy chain ratios (1:1, 1:2, 1:4, and 1:8) were tested to evaluate optimal antibody assembly. After 48 h incubation, cell culture supernatants were collected, and trastuzumab concentrations were quantified using a trastuzumab-specific ELISA kit (GenScript, Cat. No. L00970) according to the manufacturer’s instructions. Results were expressed as ng/mL of secreted antibody.

### Antibody-dependent cellular cytotoxicity (ADCC)-like assay

To assess the functional activity of secreted trastuzumab, RAW264.7 macrophage cells were seeded at 2 × 10⁴ cells per well. After 24 h, cells were treated with serum (20 μL per well) obtained from mice injected with HHES LNPs encapsulating trastuzumab mRNA (2 mg/kg, collected at day 1 post-injection). Following incubation, RAW264.7 cells were co-cultured with BT-474-luc2 target cells (HER2-positive) under standard conditions. After 24 h, target cell viability was evaluated by luminescence measurement, and cytotoxicity was expressed as relative luminescence compared to control groups.

### *In Vivo* Expression of Trastuzumab

Female C57BL/6N mice (7 weeks old) were intravenously injected with LNPs encapsulating trastuzumab heavy and light chain mRNAs at a 2:1 weight ratio. The administered dose was 2 mg/kg of total mRNA. Blood samples were collected at 1, 2, and 7 days post-injection via cheek bleeding under isoflurane anesthesia. Plasma was separated by centrifugation at 2,000 × g for 10 min at 4 °C. Trastuzumab concentrations in plasma were quantified using a trastuzumab-specific ELISA kit (GenScript, Cat. No. L00970) according to the manufacturer’s protocol. Results were expressed as ng/mL of antibody in plasma.

### Cell-Specific Delivery of LNPs in Liver and Lung Using LSL-tdTomato Mice

All animal procedures were approved by the Institutional Animal Care and Use Committee of Ewha Womans University. LSL-tdTomato mice (strain 007914, The Jackson Laboratory) were intravenously injected with Cre mRNA-loaded LNPs at a dose of 0.01 mg/kg at 8 weeks of age. After 48 h, mice were anesthetized with isoflurane and perfused with 25 mL PBS (2% FBS, 1% penicillin/streptomycin; Cytiva, SH30028.02). Perfused livers were minced in 2 mL RPMI (Gibco, 21875034) and digested in a 5 mL tube containing 2 mL RPMI supplemented with Liberase (100 μL, Roche, Cat. No. 5401127001), DNase I (5 μL, Roche, Cat. No. 4716728001), and hyaluronidase (100 μL, Sigma-Aldrich, H3506). Tissues were incubated at 37 °C, 200 rpm for 30–60 min, and then filtered through a 70-μm nylon mesh strainer (BD Falcon, Cat. No. 93070). The single-cell suspension was washed with PBS (2% FBS, 1% penicillin/streptomycin), centrifuged at 400–500 × g for 5 min, and resuspended in cell staining buffer (BioLegend, Cat. No. 420201). Cells were pre-incubated with TruStain FcX™ PLUS (BioLegend, Cat. No. 156603) for Fc receptor blocking, followed by staining with APC-anti-CD31, FITC-anti-CD45, and PE/Cy7-anti-CD68 (BioLegend) in the dark for 15 min at 4 °C. Stained cells were washed, resuspended, and analyzed using a NovoCyte 2060R flow cytometer (ACEA Biosciences). A minimum of 1.5 × 10⁶ events were collected per sample for subsequent gating and compensation analysis.

### *In Vivo* TTR Knockout Using Cas9 mRNA/sgRNA LNPs sgRNA design and chemistry

The target sequence within mouse Ttr was 5′-TTACAGCCACGTCTACAGCA-3′. The sgRNA was chemically stabilized with 2′-O-methyl and phosphorothioate modifications at the first three 5′ and 3′ terminal RNA residues and supplied as dry powder (Genscript, Piscataway, NJ, USA). Full sgRNA sequences are provided in **Supplementary Table 2a**.

### Formulation and dosing

Cas9 mRNA and sgRNA were co-formulated into LNPs (HHES or LP-01) at a 1:1 (w/w) mRNA:sgRNA ratio. Female C57BL/6N mice (7 weeks old) were administered a single intravenous injection via the retro-orbital vein at 2.0 mg/kg total RNA. For dose–response experiments, an additional low-dose cohort (1.0 mg/kg) was included.

### Plasma TTR quantification

Blood was collected at baseline (pre-dose) and at 1, 2, and 7 days post-injection for acute profiling, and weekly up to 4 weeks for durability. Plasma was separated by centrifugation (2,000 × g, 10 min, 4 °C). TTR protein levels were quantified using a mouse TTR ELISA kit (MyBioSource, Cat. No. MBS2022393) according to the manufacturer’s instructions. Results were expressed as TTR percentage (%) normalized to individual pre-dose baseline values.

### Indel analysis

At the terminal time point, livers were harvested and genomic DNA was extracted. A ∼250–300 bp amplicon spanning the *Ttr* target site was PCR-amplified and sequenced. Editing efficiency (indel %) was quantified by targeted amplicon next-generation sequencing and analyzed using CRISPResso2.

### *In Vivo* Subretinal Injection of mRNA / LNPs

All animal procedures involving ocular experiments were conducted in accordance with the guidelines of the Association for Research in Vision and Ophthalmology (ARVO) Statement for the Use of Animals in Ophthalmic and Vision Research and were approved by the IACUC of Yonsei University (IACUC-2023-0245). Mice were housed under standard laboratory conditions with a 12-hour light/dark cycle and ad libitum access to food and water.

For subretinal injections, mice were anesthetized by intraperitoneal administration of tiletamine/zolazepam combined with xylazine, and pupils were dilated with 1% tropicamide. A small incision was made at the corneal limbus using a 30-gauge needle. A Nanofil gas-tight injection system (World Precision Instruments, Sarasota, FL, USA) fitted with a 36-gauge blunt needle was introduced through the limbal incision and advanced through the vitreous cavity into the subretinal space under direct visualization using a surgical microscope.

All LNP formulations were administered by subretinal injection in a total volume of 1 µL per eye. The suspension was delivered slowly to generate a localized subretinal bleb. The needle was withdrawn carefully to minimize reflux of the injected solution, and the fundus was immediately examined to confirm successful bleb formation. Eyes exhibiting vitreous hemorrhage or other overt surgical complications were excluded from subsequent analyses.

### Assessment of Retinal Toxicity of LNPs

To evaluate the retinal toxicity of LNP formulations, fLuc mRNA-loaded LNPs formulated with HHES, MC3, or SM-102 were administered by subretinal injection to C57BL/6J mice (8 weeks old) at concentrations of 200 or 400 ng/µL (total RNA dose per eye: 200 or 400 ng in 1 µL). To comprehensively evaluate retinal toxicity, both structural and functional assessments were performed 3 weeks after injection. Structural analyses included histological evaluation of retinal architecture by hematoxylin and eosin (H&E) staining, assessment of microglial activation by IBA1 immunofluorescence, and detection of apoptotic cell death by TUNEL assay. Retinal function was assessed by full-field electroretinography (ERG).

### H&E Staining

Eyes were enucleated and fixed in Hartman’s Fixative (Sigma-Aldrich, H0290) for 40 h at room temperature. Fixed tissues were processed through graded ethanol, cleared in xylene, and embedded in paraffin. Sections were cut at 4–5 µm and stained with H&E. Bright-field images were acquired using a light microscope (BX54F; Olympus, Tokyo, Japan). Retinal thickness and outer nuclear layer (ONL) thickness were measured using ImageJ software (NIH). Measurements were performed on histological sections in which the needle track through the neural retina was identifiable. To avoid confounding by mechanical damage at the injection site, measurements were taken at a location 300–500 µm from the needle entry site in the direction of the optic nerve head.

### IBA1 Immunofluorescence Staining

Paraffin-embedded retinal sections were deparaffinized, rehydrated, and subjected to heat-mediated antigen retrieval using Target Retrieval Solution, Citrate pH 6 (Agilent Dako, Santa Clara, CA, USA; S236984-2) at 95–100 °C for 20 min. Sections were blocked in 2% normal donkey serum (NDS), 5% bovine serum albumin (BSA), and 0.3% Triton X-100 in PBS for 1 h at room temperature, followed by overnight incubation at 4 °C with a primary antibody against IBA1 (1:1000; Abcam, Cambridge, UK; ab178846). After washing, sections were incubated with Alexa Fluor 488-conjugated donkey anti-rabbit IgG (1:2000; Invitrogen, Carlsbad, CA, USA; A-21206) for 1 h at room temperature. Sections were mounted using VECTASHIELD Antifade Mounting Medium containing DAPI (Vector Laboratories, Newark, CA, USA; H-1200) for nuclear counterstaining. Fluorescence images were acquired using an inverted fluorescence microscope (IX73-F22PH; Olympus, Tokyo, Japan). IBA1-positive area normalized to DAPI-positive area was quantified using ImageJ software (NIH).

### TUNEL Assay

Apoptotic cells were detected using the In Situ Cell Death Detection Kit, Fluorescein (Roche Diagnostics, 11684795910) according to the manufacturer’s instructions. Paraffin-embedded retinal sections were deparaffinized, rehydrated, and subjected to antigen retrieval as described above. Sections were incubated with the TUNEL reaction mixture for 1 h at room temperature in the dark. Sections were mounted with VECTASHIELD containing DAPI as described above. Fluorescence images were acquired using an inverted fluorescence microscope (IX73-F22PH). The percentage of TUNEL-positive cells relative to DAPI-positive nuclei was quantified using ImageJ software (NIH).

### Electroretinography (ERG)

ERG was performed using the Celeris Electroretinography system (Diagnosys LLC, Lowell, MA, USA) to assess retinal function. Mice were dark-adapted overnight (>12 h) prior to recordings. Following anesthesia by intraperitoneal injection of tiletamine/zolazepam and xylazine, pupils were dilated with 1% tropicamide. The corneal surface was anesthetized with 0.5% proparacaine hydrochloride to minimize discomfort and motion artifacts. Body temperature was maintained at 37 °C using a thermostatically controlled heating pad.

A reference electrode was positioned in the oral cavity, a ground electrode was placed subcutaneously at the base of the tail, and active corneal electrodes were gently positioned on the corneal surface with a layer of methylcellulose to ensure optimal electrical contact. Scotopic (dark-adapted) ERG responses were recorded using a series of increasing flash stimuli from −2.5 to 1.5 log cd·s/m². Photopic (light-adapted) responses were recorded at 0.5 and 1.0 log cd·s/m² following a 10-minute light adaptation period at a background luminance of 9 cd/m². After recordings, mice were monitored until full recovery from anesthesia and administered antibiotic ophthalmic ointment bilaterally. ERG waveforms were recorded and analyzed using the system’s proprietary software (Espion, Diagnosys).

### Retinal Delivery of LNPs Using LSL-tdTomato Mice

LSL-tdTomato reporter mice (Ai9; strain 007909, The Jackson Laboratory), in which Cre-mediated excision of a loxP-flanked STOP cassette induces tdTomato expression, received subretinal injections of HHES-Cre mRNA-loaded LNPs (400 ng/µL) as described above. Reporter activation was evaluated 1 weeks after injection by whole-mount imaging of the RPE/choroid complex and retinal cryosections. For whole-mount preparation, mice were euthanized and eyes were enucleated and rinsed in PBS. Eyeballs were fixed in 1% paraformaldehyde (PFA) in PBS for 1 h. The anterior segment, including the cornea, iris, and lens, was removed in PBS. The neural retina was carefully separated, leaving the RPE/choroid complex, which was radially incised into eight segments centered on the optic disc. Tissues were flat-mounted on glass slides and mounted using VECTASHIELD Antifade Mounting Medium containing DAPI for nuclear counterstaining. For retinal cryosections, enucleated eyes were rinsed with PBS and the anterior segment was removed as described above. Samples were fixed in 4% paraformaldehyde containing 5% sucrose in PBS for 30 min, followed by sequential cryoprotection in 10%, 20%, and 30% sucrose solutions until equilibrated at each concentration. Tissues were embedded in OCT compound, frozen, and sectioned at 7 µm using a cryostat. Sections were mounted using VECTASHIELD Antifade Mounting Medium containing DAPI. Fluorescence images were acquired using an inverted fluorescence microscope (IX73-F22PH; Olympus, Tokyo, Japan).

### Retinal Delivery of LNPs in Human Retinal Organoids and RPE cells

Human retinal organoids were differentiated from human embryonic stem cells (hESCs; H9) following a previously described protocol with minor modifications [39]. Mature retinal organoids cultured for more than 30 weeks were used for experiments. RPE cells were generated from H9 hESCs using a previously reported differentiation method and maintained in culture until they exhibited characteristic RPE morphology, including pigmentation and hexagonal organization [40]. RPE identity was further confirmed by immunostaining for RPE65 and ZO-1.

For mRNA delivery experiments, HHES LNPs encapsulating GFP mRNA were applied to retinal organoids or RPE cells. Retinal organoids were treated with HHES-GFP mRNA LNPs at a concentration of 10 µg /mL, whereas RPE cells were treated with 0.25 µg/mL or 1 µg/mL HHES-GFP mRNA LNPs. After 24 h, GFP expression was evaluated by live fluorescence imaging. Brightfield and fluorescence images were acquired using an inverted fluorescence microscope (IX73-F22PH).

### Design and *in vitro* validation of *Vegfa*-targeting sgRNAs

For SpCas9-mediated genome editing, a previously reported sgRNA targeting the mouse *Vegfa* locus (5′-CTCCTGGAAGATGTCCACCA-3′) was used [41]. For ABE8e-mediated base editing, an sgRNA targeting the *Vegfa* intron 1 splice donor site (5′-CGCTTACCTTGGCATGGTGG-3′) was selected using SpliceR v1.3.0 [42], with the base-editing window set to positions 4–8. This sgRNA was predicted to disrupt the splice donor and was assigned a high ABE score (0.84), a sequence-context- and target-position-based ranking metric for predicted ABE editing. An sgRNA targeting the *Rosa26* locus (5′-GGCGGTCCTCAGAAGCCAGG-3′) was used as a non-targeting control.

sgRNA sequences were cloned into the BsaI-digested pRG2 vector (#104174, Addgene, Watertown, MA, USA), and all constructs were verified by Sanger sequencing. Mouse Neuro-2a cells (ATCC CCL-131, American Type Culture Collection, Manassas, VA, USA) were transfected with pRGEN-Cas9-CMV/T7-Puro-RFP (ToolGen, Seoul, Korea) or ABE8e expression plasmid (#138489, Addgene), together with the corresponding sgRNA expression vectors, using Lipofectamine 2000 (Thermo Fisher Scientific; 11668027) according to the manufacturer’s instructions. Cells were harvested 96 hours post-transfection for downstream analyses.

To validate the reduction of VEGFA protein, western blot analysis was performed using anti-VEGFA (Abcam; ab214424) and anti-β-actin (Santa Cruz Biotechnology, Dallas, TX, USA; sc-47778) antibodies. Following incubation with HRP-conjugated anti-rabbit (7074) or anti-mouse (7076) secondary antibodies (Cell Signaling Technology, Danvers, MA, USA), signals were detected using ECL solution (Bio-Rad Laboratories, CA, USA). Images were acquired using an Amersham ImageQuant 800 (Cytiva, Tokyo, Japan).

### *In Vivo* Retinal Genome Editing sgRNA preparation

All sgRNAs were chemically synthesized with 2′-O-methyl and phosphorothioate modifications at the first three and last three nucleotides (Bionics, Seoul, Korea). Lyophilized sgRNAs (10 nmol) were reconstituted according to the manufacturer’s instructions. The full modified sequences are listed in **Supplementary Table 2a**.

### Formulation and dosing

SpCas9 mRNA and sgRNA were co-formulated into HHES LNPs at a 1:1 (w/w) mRNA:sgRNA ratio. ABE8e mRNA and sgRNA were also co-formulated similarly. C57BL/6J mice (8 weeks old) received subretinal injections of the indicated HHES LNP formulations at a total RNA concentration of 300 ng/µL (150 ng/µL mRNA + 150 ng/µL sgRNA), as described above. For genome-editing efficiency analysis, eyes were enucleated two weeks after injection (n = 3 eyes for *Rosa26* sgRNA and n = 5 eyes for *Vegfa* sgRNA in the SpCas9 experiment; n = 3 eyes per group in the ABE8e experiment). After removal of the anterior segment and vitreous, the eyecup was quartered, and the neural retina was carefully removed. The remaining RPE–choroid–sclera complex was incubated in 0.25% trypsin–EDTA at 37 °C for 45 min, washed with PBS, and gently tapped to dissociate RPE cells from the underlying choroid. Released RPE cells were collected by centrifugation (300 × g, 4 °C, 5 min), and genomic DNA was extracted for targeted deep sequencing.

### Genomic DNA extraction

Genomic DNA was extracted by adding 20 μL of freshly prepared lysis buffer (10 mM Tris-HCl, pH 8.0, 0.05% SDS, and 25 μg/mL proteinase K; Thermo Fisher Scientific) directly to each sample. Samples were incubated at 37 °C for 1 h. The lysates were transferred to a 96-well PCR plate and incubated at 80 °C for 15 min to inactivate proteinase K.

### Targeted deep sequencing

Target regions were amplified from the extracted genomic DNA using locus-specific primers (**Supplementary Table 2b**). Deep-sequencing libraries were generated by PCR, and TruSeq HT dual-index primers were used to label individual samples. Pooled libraries were subjected to paired-end sequencing using HiSeq X (Illumina, San Diego, CA, USA). Indel frequencies for SpCas9 were quantified using Cas-Analyzer [43], and on-target A-to-G editing efficiencies for ABE8e were quantified using BE-Analyzer [44].

### Laser-Induced Choroidal Neovascularization (CNV) Model

In a separate cohort of C57BL/6J mice (8 weeks old), laser-induced CNV was generated 2 weeks after subretinal injection of HHES LNPs containing the indicated genome editing components. CNV lesions were analyzed 1 week after laser induction. CNV was induced by laser photocoagulation. Following anesthesia by intraperitoneal administration of tiletamine/zolazepam and xylazine and pupil dilation with 1% tropicamide, three laser burns were applied to each eye using a 532 nm laser photocoagulation system (GYC-1000; NIDEK CO., LTD., Gamagori, Aichi, Japan) with the following parameters: 50 µm spot size, 100 mW power, and 50 ms exposure time. Laser spots were placed approximately two disc diameters from the optic nerve head and evenly distributed around the posterior pole. Only burns that produced a cavitation bubble, indicating rupture of Bruch’s membrane, were included in the analysis. Eyes with subretinal hemorrhage were excluded.

For choroidal flat-mount preparation, eyes were enucleated 1 week after laser induction and fixed in 4% paraformaldehyde in PBS for 1 h at room temperature. The anterior segment and neural retina were removed, and the RPE/choroid/sclera complex was isolated. Tissues were permeabilized and stained with Isolectin GS-IB4 conjugated to Alexa Fluor 488 (1:100; Invitrogen, Carlsbad, CA, USA; I21411) in PBS containing 0.3% Triton X-100 overnight at 4 °C. After washing with PBS, flat mounts were prepared by making radial incisions and mounted on glass slides for imaging.

CNV lesion areas were quantified using ImageJ software (NIH). The neovascular area of each lesion was measured by a masked observer. Three laser-induced lesions were generated per eye, and all qualifying lesions were included in the quantitative analysis. For the SpCas9 experiment, a total of 18 qualifying lesions from 6 eyes were analyzed per group; for the ABE8e experiment, 12 qualifying lesions from 4 eyes were analyzed per group.

## Statistical Analysis

Data are presented as mean ± standard error of the mean (SEM). Retinal thickness, ONL thickness, IBA1-positive area, and TUNEL-positive cells were analyzed using one-way ANOVA followed by Bonferroni’s multiple-comparisons test. Neovascular surface area between *Rosa26* control and *Vegfa*-targeted groups was compared using an unpaired two-tailed Student’s t-test. P < 0.05 was considered statistically significant. All statistical analyses were performed using GraphPad Prism (version 10.6.1; GraphPad Software, San Diego, CA, USA).

## Use of Artificial Intelligence

Manuscript drafting was assisted by Claude (Anthropic), an AI language model. All AI-generated content was critically reviewed, edited, and verified by the authors, who take full responsibility for the accuracy and integrity of the reported findings

## Acknowledgements

This study was supported by a grant from the Korea Institute of Radiological and Medical Sciences (KIRAMS), Ministry of Science and ICT (MSIT), Republic of Korea (No. 50461-2026). This work was also supported by the Bio & Medical Technology Development Program of the National Research Foundation of Korea (NRF), funded by the Ministry of Science and ICT (MSIT) (grant numbers: RS-2023-00261343, RS-2024-00451880, RS-2024-00411768, RS-2025-00542986, RS-2024-00465298, and RS-2025-25455885), and by the National Research Foundation of Korea (NRF) grant funded by the Korean government (MSIT) (grant number: RS-2025-00514263). This research was also supported by the Korea Health Technology R&D Project through the Korea Health Industry Development Institute (KHIDI), funded by the Ministry of Health and Welfare, Republic of Korea (grant number: RS-2025-25459112). Schematic figures were created with BioRender.com.

