## Supplementary Data for "A Rapidly Excretable, ROS-Scavenging Ionizable Lipid Decouples mRNA Delivery Potency from Toxicity"

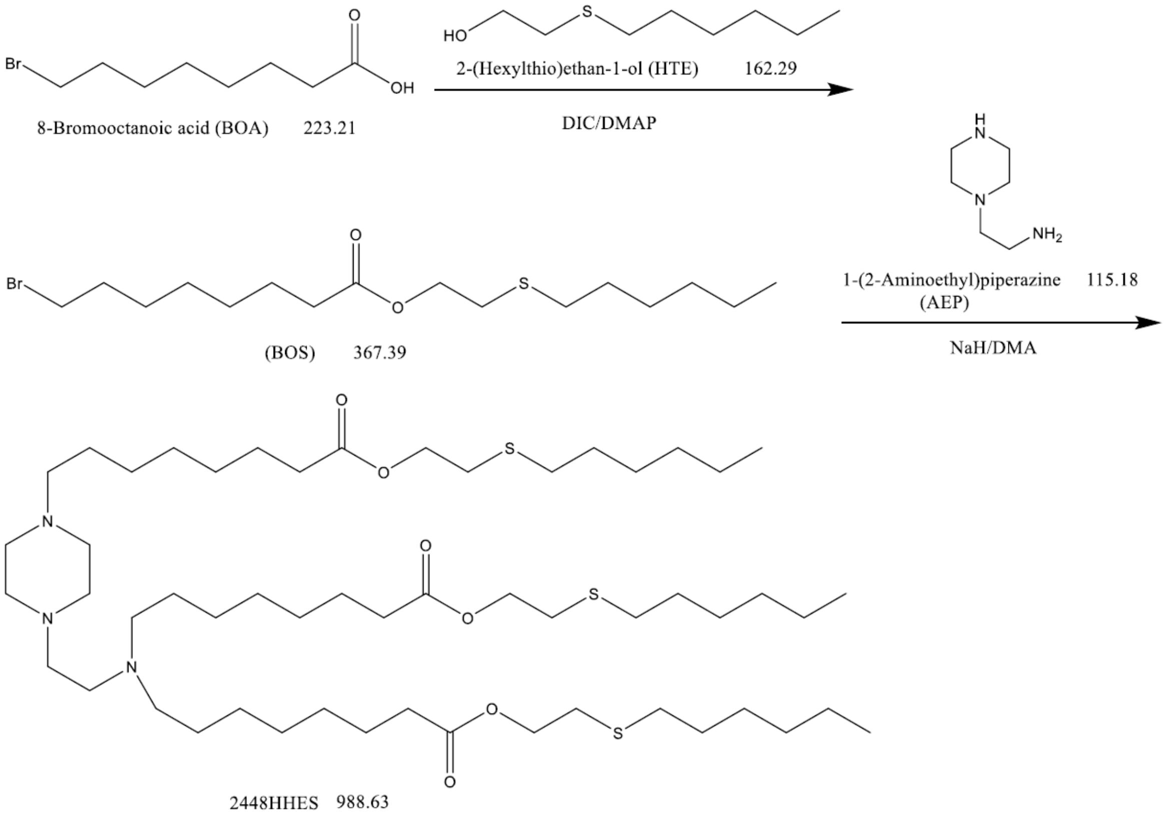


**Supplementary Figure 1. Synthesis of b-3-HHES.** 8-Bromooctanoic acid (BOA) was coupled with 2-(hexylthio)ethan-1-ol (HTE) in the presence of DIC and DMAP in DCM, producing the intermediate BOS. BOS was then reacted with 1-(2-aminoethyl)piperazine (AEP) using NaH in DMA to yield the ionizable lipid 2448HHES. The final product was purified by column chromatography. *m/z* calcd for $C_{52}H_{104}N_3O_6S_2^+$ (M+H), 988.63; found, 988.63.

**Compound Table**

| Compound Label | RT | Mass | Abund | Formula | Tgt Mass | Diff (ppm) |
|---|---|---|---|---|---|---|
| Cpd 1: C54 H105 N3 O6 S3 | 0.188 | 987.7179 | 10777 | C54 H105 N3 O6 S3 | 987.7166 | 1.41 |

---

| Compound Label | *m/z* | RT | Algorithm | Mass |
|---|---|---|---|---|
| Cpd 1: C54 H105 N3 O6 S3 | 1010.7056 | 0.188 | Find By Formula | 987.7179 |

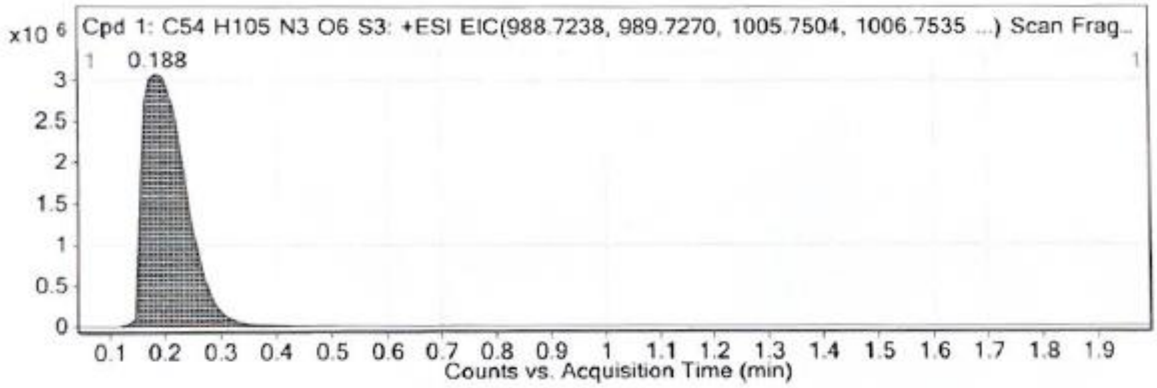


MS Spectrum

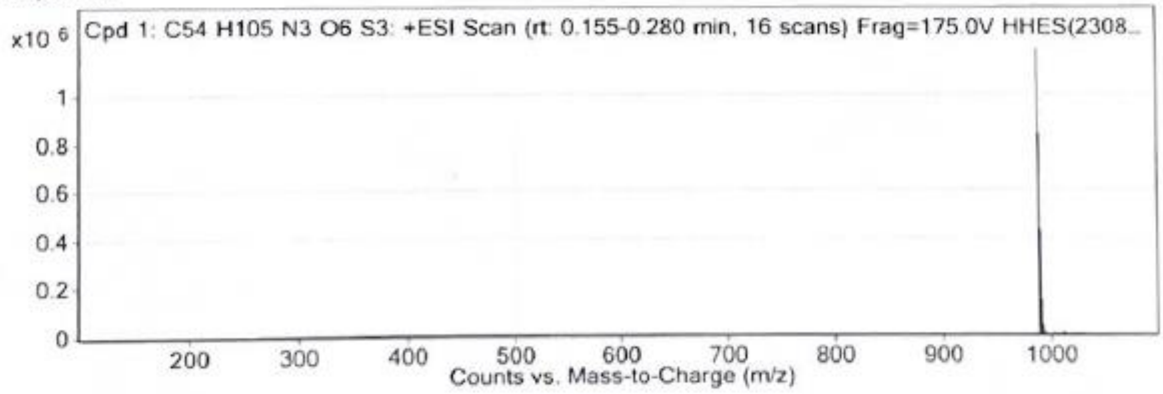


MS Zoomed Spectrum

**Supplementary Figure 2. LC/MS-TOF analysis of b-3-HHES.** LC/MS-TOF analysis confirmed the successful formation of b-3-HHES. The extracted ion chromatogram (EIC) showed a single dominant peak at RT = 0.188 min, consistent with the expected molecular ion. The corresponding mass spectrum exhibited a major ion at m/z 987.7179, which matches the calculated mass for the proposed formula ($C_{54}H_{105}N_3O_6S_3^+$, calcd 987.7166), with a mass error of 1.41 ppm. These results verify the high purity and correct molecular composition of b-3-HHES.

**Compound Table**

| Compound Label | RT | Mass | Abund | Formula | Tgt Mass | Diff (ppm) |
|---|---|---|---|---|---|---|
| Cpd 1: C56 H111 N3 O6 S3 | 0.384 | 1017.7625 | 1618 | C56 H111 N3 O6 S3 | 1017.7635 | -0.98 |

| Compound Label | *m/z* | RT | Algorithm | Mass |
|---|---|---|---|---|
| Cpd 1: C56 H111 N3 O6 S3 | 1018.7702 | 0.384 | Find By Formula | 1017.7625 |

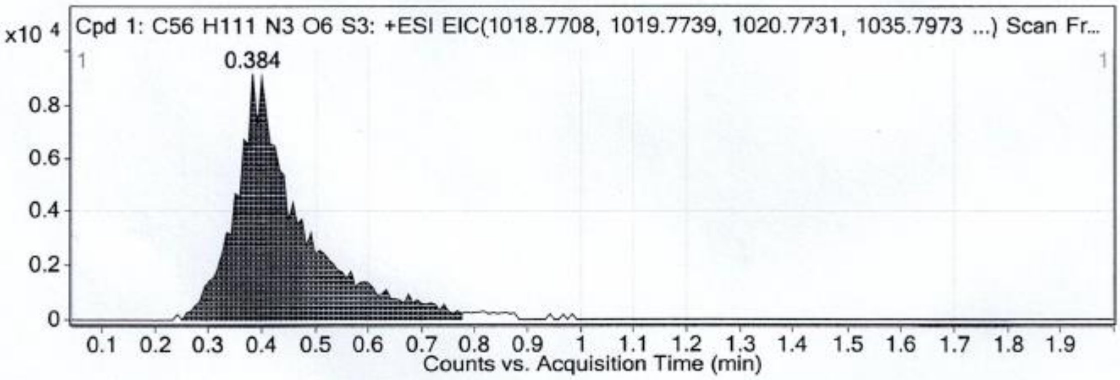


MS Spectrum

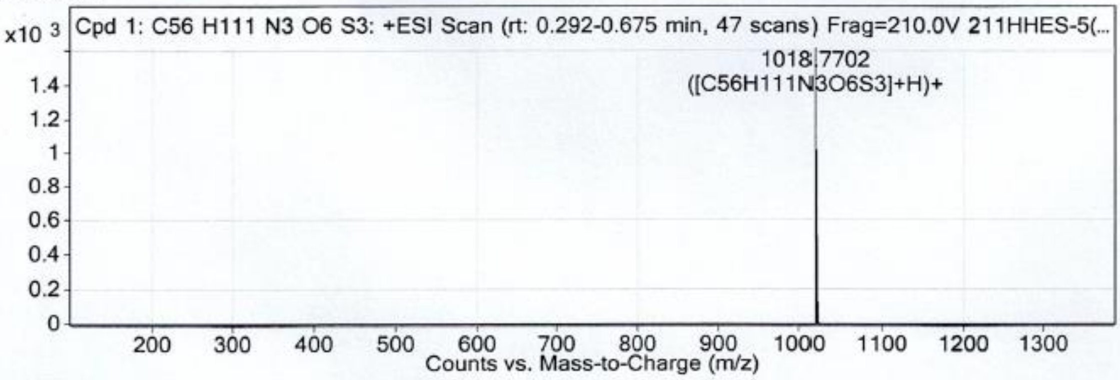


MS Zoomed Spectrum

**Supplementary Figure 3. LC/MS-TOF analysis of a-3-HHES.** LC/MS-TOF analysis of a-3-HHES showed a clear chromatographic peak at RT = 0.384 min, indicating a well-defined single component. The mass spectrum displayed a dominant ion at m/z 1017.7625, in excellent agreement with the calculated mass for the proposed molecular formula ($C_{56}H_{111}N_3O_6S_3^+$, calcd 1017.7635; Δ = –0.98 ppm). The agreement between the observed and theoretical mass values confirms the correct molecular composition and high purity of a-3-HHES.

**Compound Table**

| Compound Label | RT | Mass | Abund | Formula | Tgt Mass | Diff (ppm) |
|---|---|---|---|---|---|---|
| Cpd 1: C74 H144 N4 O8 S4 | 0.248 | 1344.9861 | 5220 | C74 H144 N4 O8 S4 | 1344.9867 | -0.41 |

| Compound Label | *m/z* | RT | Algorithm | Mass |
|---|---|---|---|---|
| Cpd 1: C74 H144 N4 O8 S4 | 1345.9936 | 0.248 | Find By Formula | 1344.9861 |

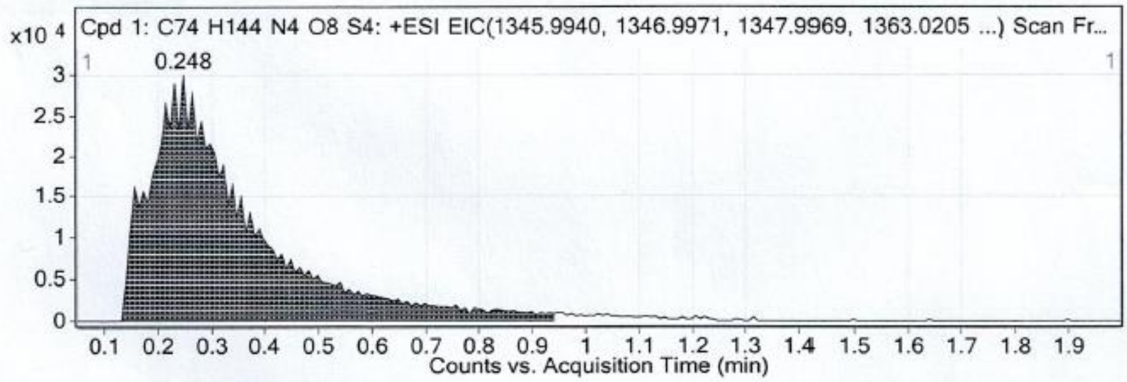


MS Spectrum

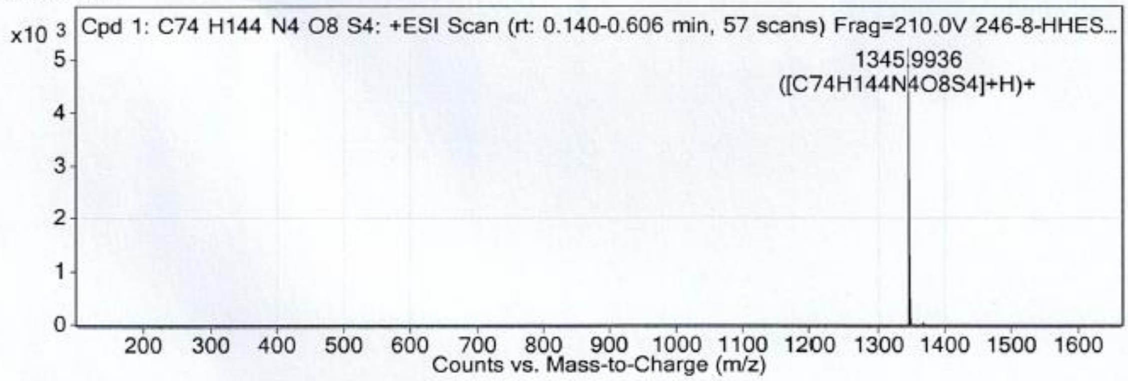


MS Zoomed Spectrum

**Supplementary Figure 4. LC/MS-TOF analysis of c-3-HHES.** LC/MS-TOF analysis of c-3-HHES showed a distinct chromatographic peak at RT = 0.248 min, indicating a single major component. The mass spectrum displayed a dominant ion at m/z 1345.9936, consistent with the calculated mass for the proposed formula ($C_{74}H_{144}N_4O_8S_4^+$, calcd 1344.9867; Δ = –0.41 ppm). The excellent agreement between observed and theoretical mass confirms the correct molecular composition and high purity of c-3-HHES.

**Compound Table**

| Compound Label | RT | Mass | Abund | Formula | Tgt Mass | Diff (ppm) |
|---|---|---|---|---|---|---|
| Cpd 1: C48H93N3O6S3 | 0.463 | 903.6245 | 115980 | C48H93N3O6S3 | 903.6226 | 2.06 |

| Compound Label | RT | Algorithm | Mass |
|---|---|---|---|
| Cpd 1: C48H93N3O6S3 | 0.463 | Find By Formula | 903.6245 |

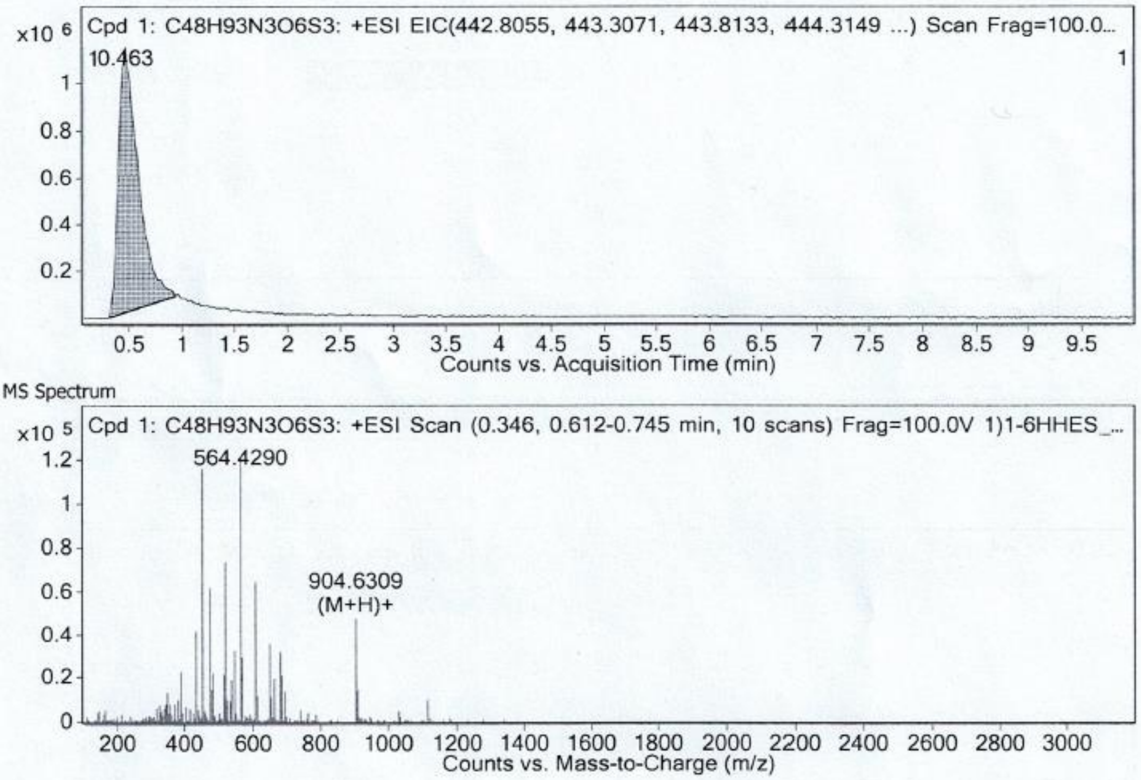


**Supplementary Figure 5. LC/MS-TOF analysis of b-1-HHES.** LC/MS-TOF analysis of b-1-HHES showed a clear chromatographic peak at RT = 0.463 min, corresponding to a single major component. The mass spectrum displayed a dominant ion at m/z 903.6245, consistent with the calculated mass for the proposed formula ($C_{48}H_{93}N_3O_6S_3^+$, calcd 903.6226; Δ = 2.06 ppm). The agreement between the observed and theoretical masses confirms the correct molecular composition and successful formation of b-1-HHES.

**Compound Table**

| Compound Label | RT | Mass | Abund | Formula | Tgt Mass | Diff (ppm) |
|---|---|---|---|---|---|---|
| Cpd 1: C51H99N3O6S3 | 0.498 | 945.6717 | 71897 | C51H99N3O6S3 | 945.6696 | 2.21 |

| Compound Label | RT | Algorithm | Mass |
|---|---|---|---|
| Cpd 1: C51H99N3O6S3 | 0.498 | Find By Formula | 945.6717 |

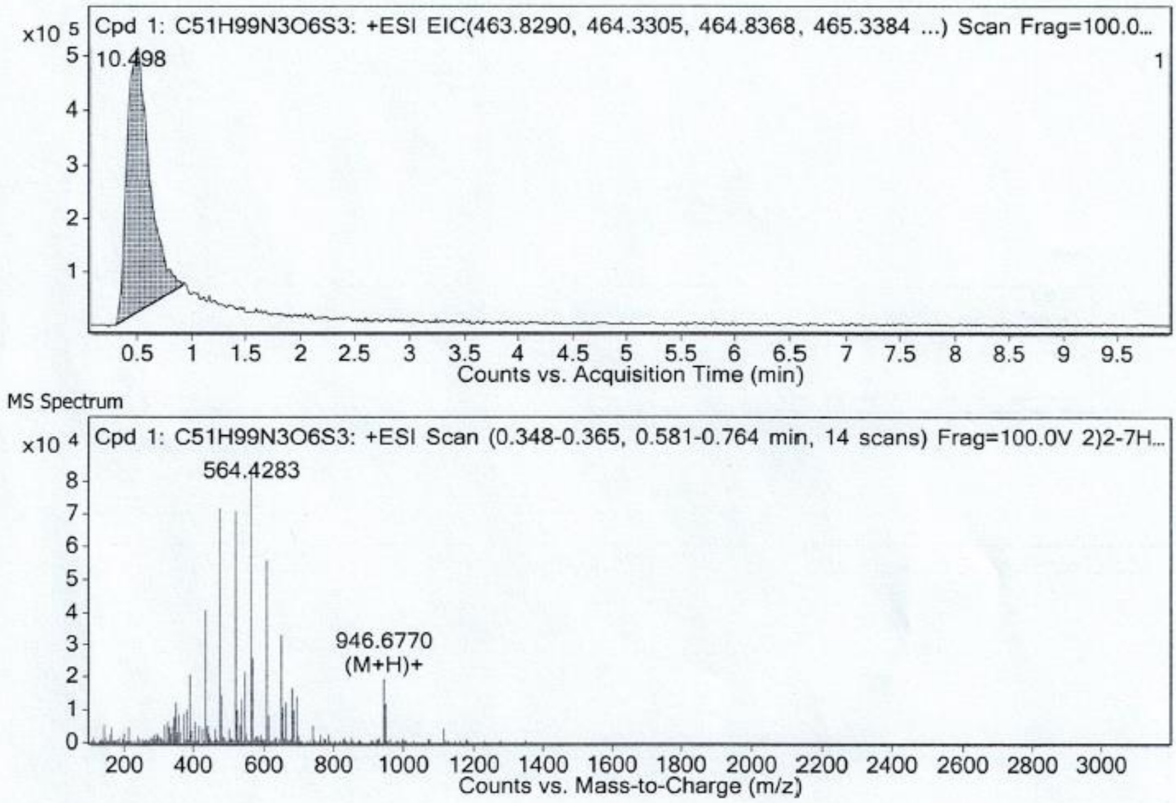


**Supplementary Figure 6. LC/MS-TOF analysis of b-2-HHES.** LC/MS-TOF analysis of b-2-HHES exhibited a distinct chromatographic peak at RT = 0.498 min, corresponding to a single major species. The mass spectrum showed a dominant ion at m/z 945.6717, consistent with the calculated mass for the proposed formula ($C_{51}H_{99}N_3O_6S_3^+$, calcd 945.6696; Δ = 2.21 ppm). The strong agreement between observed and theoretical values confirms the successful formation and correct molecular composition of b-2-HHES.

**Compound Table**

| Compound Label | RT | Mass | Abund | Formula | Tgt Mass | Diff (ppm) |
|---|---|---|---|---|---|---|
| Cpd 1: C57H111N3O6S3 | 0.547 | 1029.7654 | 107020 | C57H111N3O6S3 | 1029.7635 | 1.82 |

| Compound Label | RT | Algorithm | Mass |
|---|---|---|---|
| Cpd 1: C57H111N3O6S3 | 0.547 | Find By Formula | 1029.7654 |

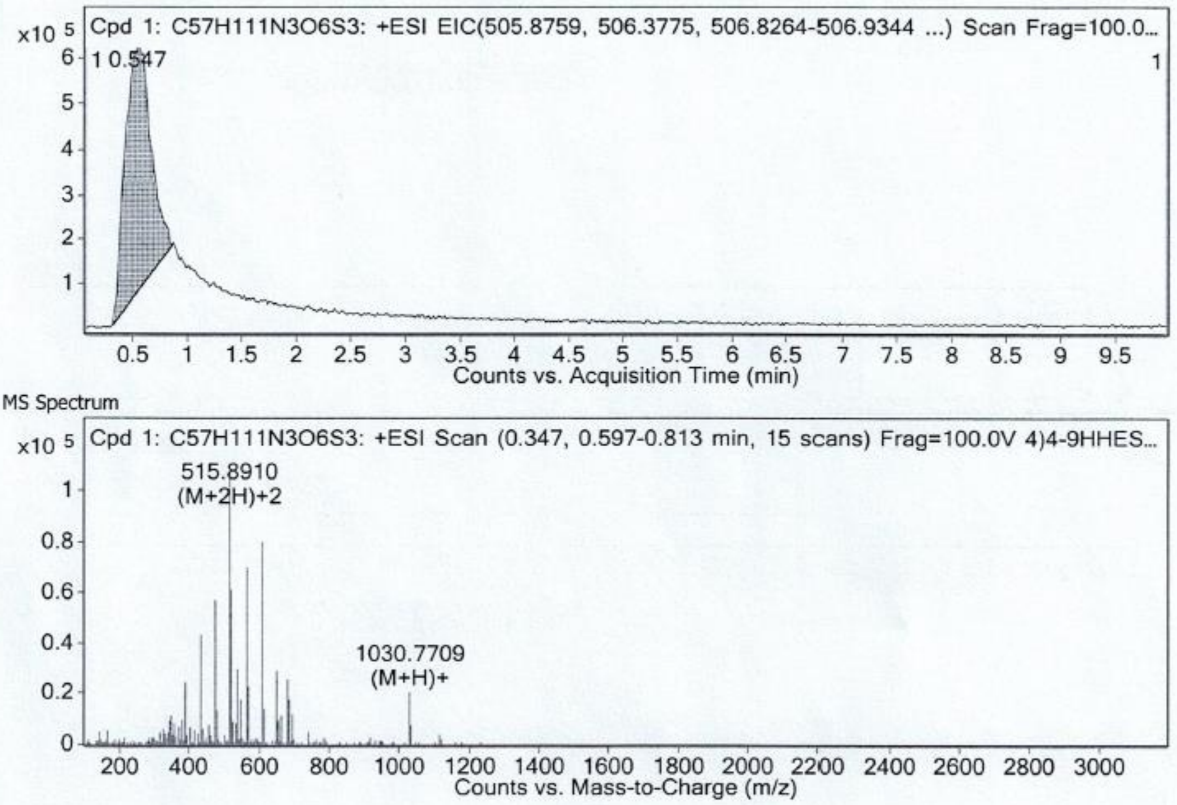


**Supplementary Figure 7. LC/MS-TOF analysis of b-4-HHES.** LC/MS-TOF analysis of b-4-HHES revealed a distinct chromatographic peak at RT = 0.547 min, indicating a single major component. The mass spectrum showed a dominant ion at m/z 1029.7654, closely matching the calculated mass for the proposed formula ($C_{57}H_{111}N_3O_6S_3^+$, calcd 1029.7635; Δ = 1.82 ppm). The high correspondence between observed and theoretical masses confirms the successful synthesis and correct molecular composition of b-4-HHES.

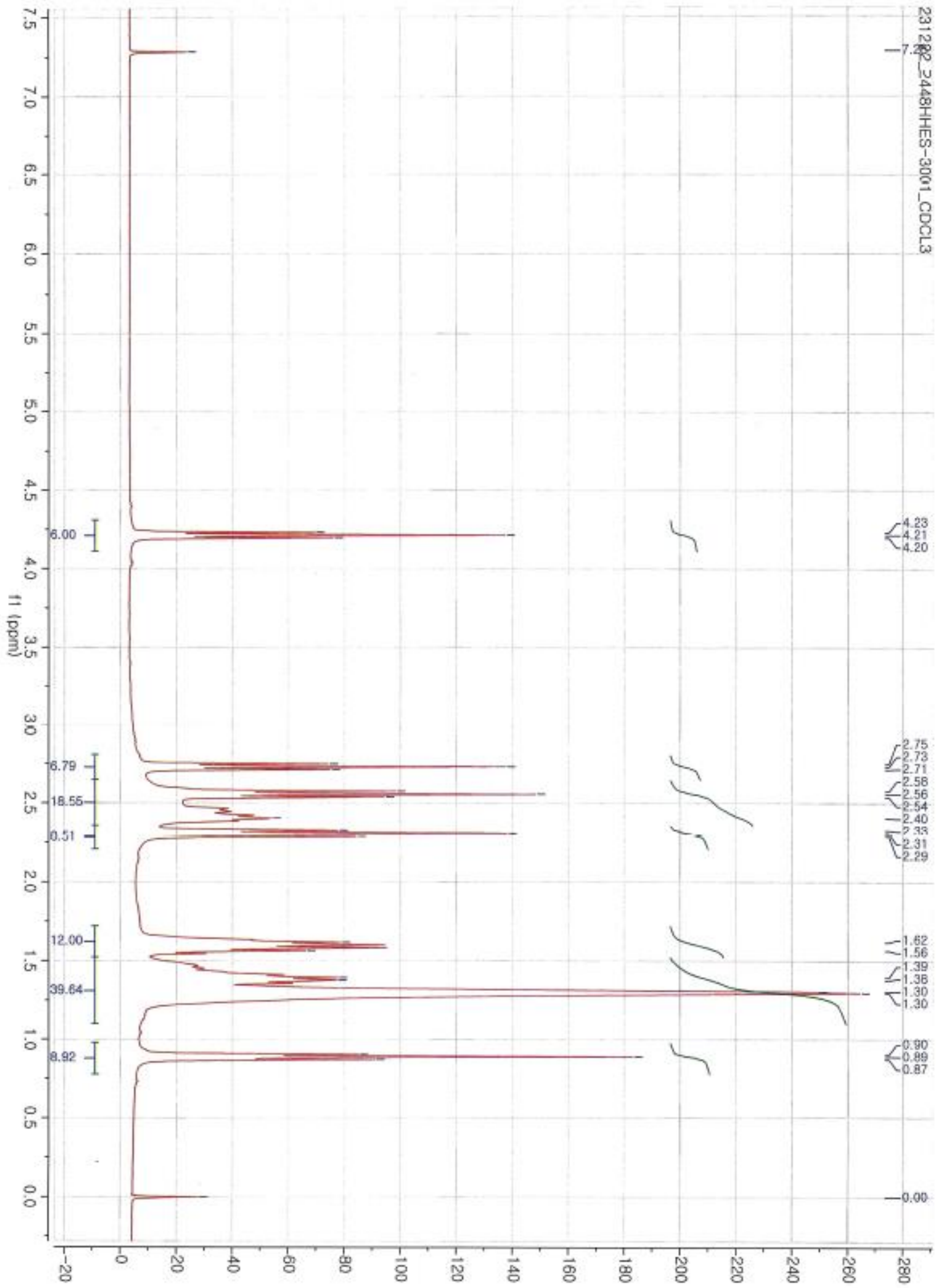

**Supplementary Figure 8. $^1$H NMR spectrum of b-3-HHES ($CDCl_3$, 300 MHz).** $^1$H NMR analysis of b-3-HHES shows characteristic proton resonances consistent with the expected molecular structure. Peaks corresponding to the aliphatic chains appear broadly between δ 0.87–1.62 ppm, while methylene protons adjacent to heteroatoms are observed at δ 2.29–2.75 ppm. Additional downfield signals assigned to protons adjacent to ester and thioether functionalities appear at δ 4.20–4.23 ppm. The overall spectral pattern confirms the successful synthesis and structural integrity of b-3-HHES.

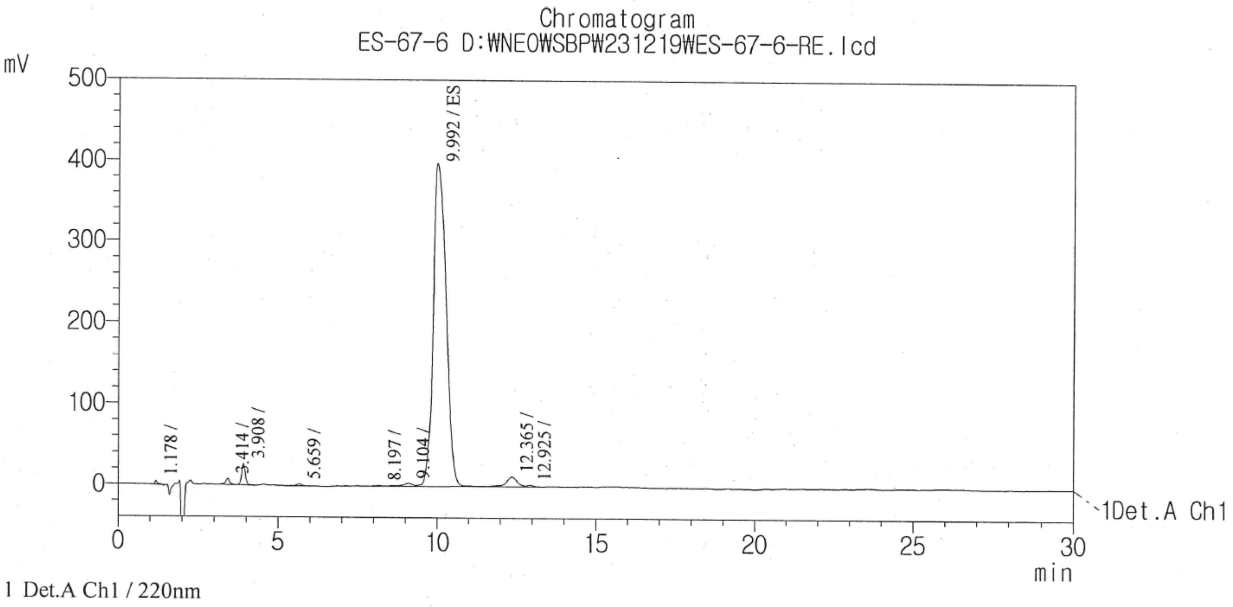


PeakTable

Detector A Ch1 220nm

| Peak# | Name | Ret. Time | Area | Area % | Height | Height % |
|---|---|---|---|---|---|---|
| 1 | | 1.178 | 24476 | 0.21 | 3517 | 0.772 |
| 2 | | 3.414 | 71850 | 0.60 | 7208 | 1.582 |
| 3 | | 3.908 | 207140 | 1.74 | 25966 | 5.699 |
| 4 | | 5.659 | 25810 | 0.22 | 2292 | 0.503 |
| 5 | | 8.197 | 10945 | 0.09 | 719 | 0.158 |
| 6 | | 9.104 | 80553 | 0.68 | 3603 | 0.791 |
| 7 | ES | 9.992 | 11170373 | 93.67 | 398267 | 87.418 |
| 8 | | 12.365 | 303091 | 2.54 | 12122 | 2.661 |
| 9 | | 12.925 | 30690 | 0.26 | 1898 | 0.417 |
| Total | | | 11924927 | 100.00 | 455592 | 100.000 |

**Supplementary Figure 9. HPLC analysis of b-3-HHES.** HPLC analysis of b-3-HHES (220 nm detection) showed a dominant peak at RT = 9.992 min, corresponding to 93.67% of the total peak area, indicating high product purity. Minor impurities were detected at earlier retention times, each contributing less than 1% of the total chromatographic area. The chromatographic profile thus confirms that b-3-HHES was obtained in high chemical purity.

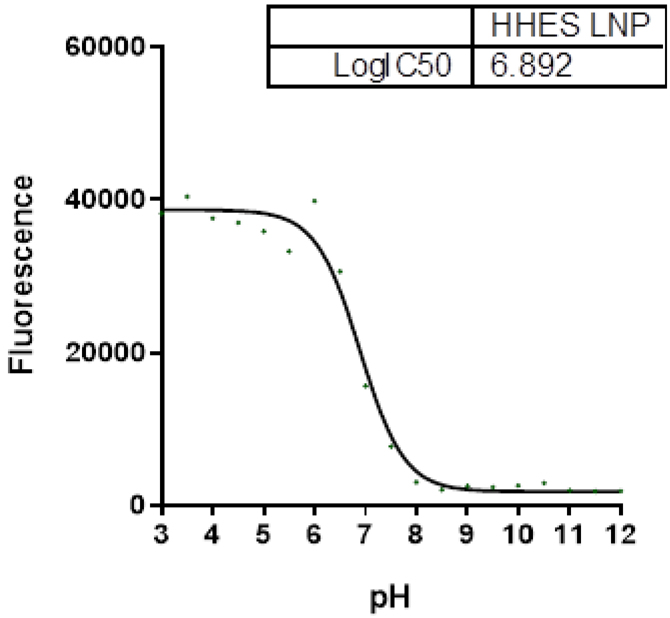


**Supplementary Figure 10. Apparent pKa of HHES LNPs measured by TNS assay.** TNS fluorescence intensity was measured across pH to determine the ionization behavior of HHES LNPs. The apparent pKa was obtained by nonlinear curve fitting (pKa = 6.81).

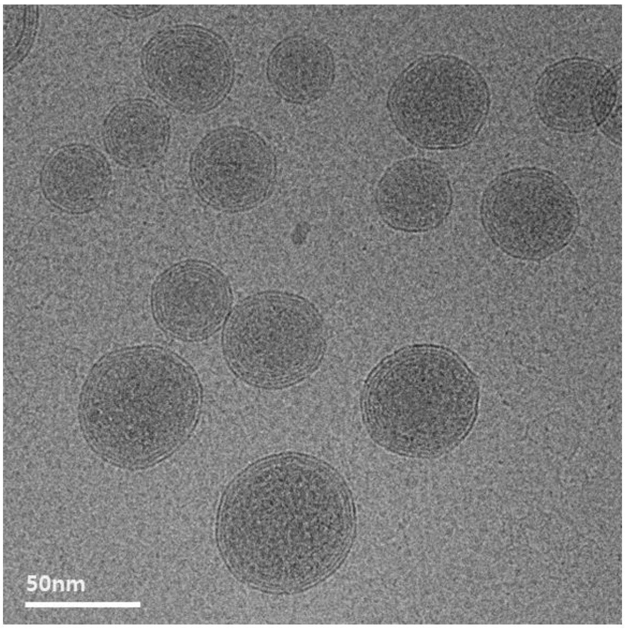


**Supplementary Figure 11. Cryogenic transmission electron microscopy (cryo-TEM) image of b-3-HHES lipid nanoparticles.** Representative cryo-TEM image showing the morphology and size distribution of b-3-HHES LNPs. Particles exhibited a predominantly spherical structure with a multilamellar internal organization. Scale bar, 50 nm.

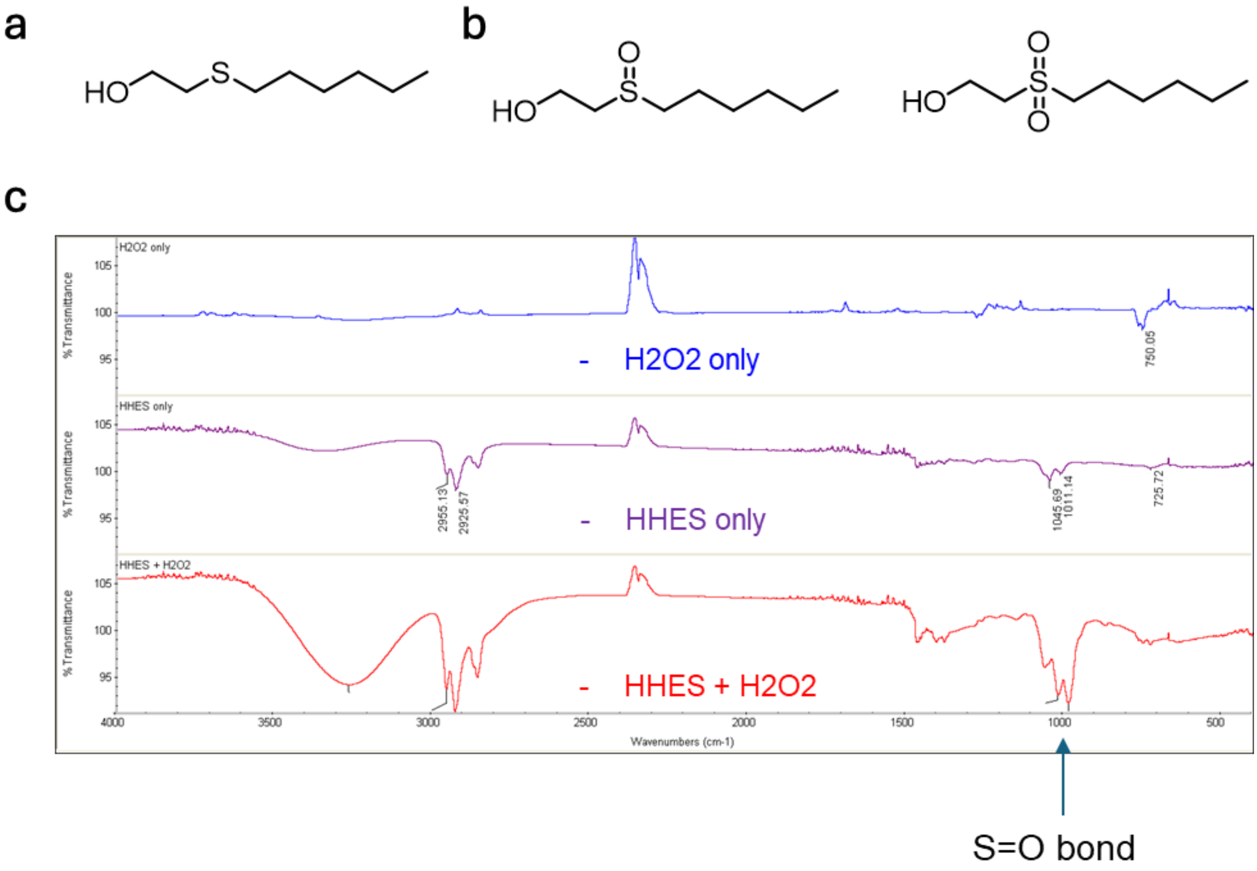


**Supplementary Figure 12. Oxidation of HHES and FT-IR confirmation of S=O formation.** (a) Chemical structure of HHES containing a thioether functional group. (b) Representative oxidized forms of HHES, illustrating possible conversion of the thioether to a sulfoxide or sulfone upon exposure to oxidizing conditions. (c) FT-IR spectra confirming oxidation of HHES. Treatment of HHES with $H_2O_2$ resulted in the appearance of a characteristic S=O stretching band at ~1000 $cm^{-1}$ (red trace), whereas $H_2O_2$ alone (blue) and untreated HHES (purple) did not exhibit this peak. These results demonstrate that HHES undergoes oxidative conversion consistent with sulfoxide/sulfone formation.

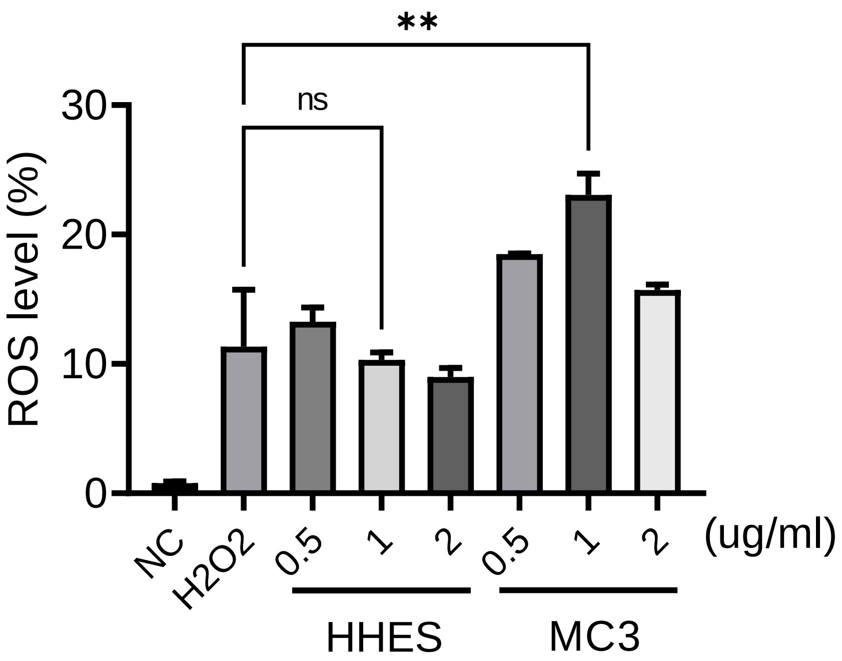


**Supplementary Figure 13. ROS scavenging activity of HHES in HEPG2 cells.** ROS levels in HEPG2 cells were measured using the DCFH-DA assay following $H_2O_2$-induced oxidative stress. Cells were treated with HHES- or MC3-based LNPs, and intracellular ROS levels were quantified by DCF fluorescence. HHES-LNPs exhibited lower ROS levels than MC3-LNPs, indicating superior ROS scavenging activity.

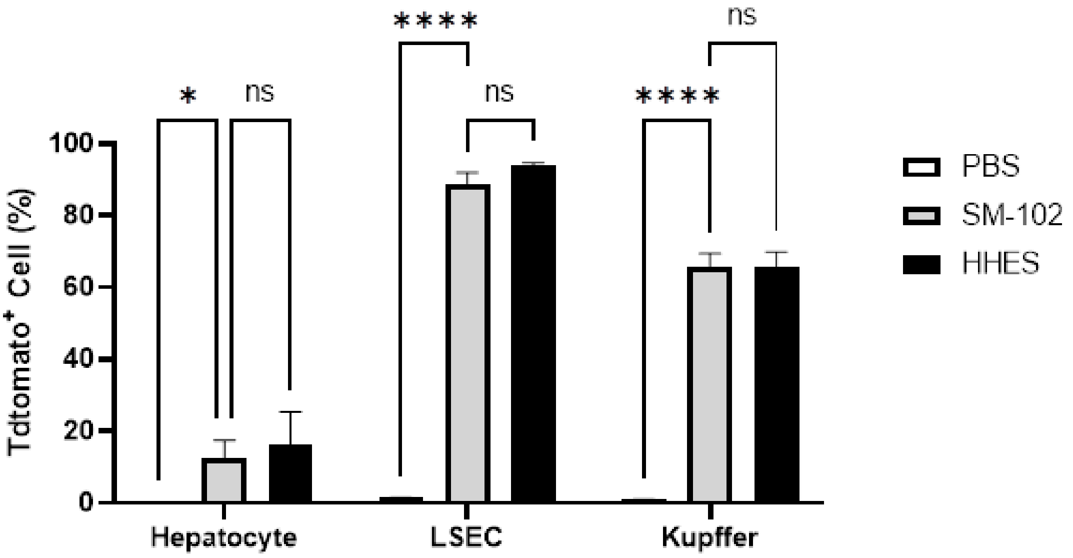


**Supplementary Figure 14. Cre-dependent tdTomato expression in liver cell types after LNP delivery in Ai14 mice.** Ai14 Cre-reporter mice were administered 0.25 mp/kg of Cre-mRNA–loaded LNPs, and liver tissues were analyzed 3 days post-injection. tdTomato expression, indicative of LNP delivery, was quantified in Kupffer cells, hepatocytes, and LSECs.

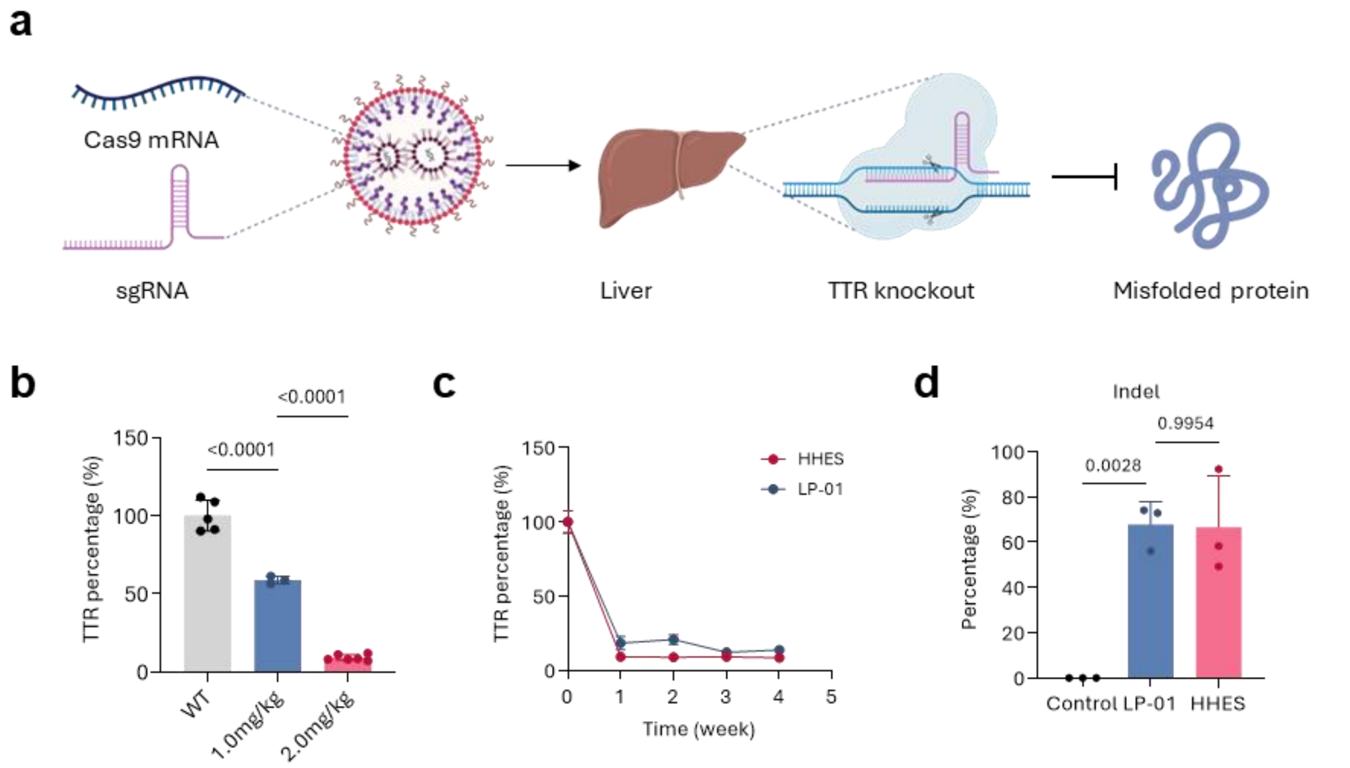


**Supplementary Figure 15. In vivo TTR knockout in mice using Cas9 mRNA–loaded LNPs.** (a) Schematic illustration of in vivo CRISPR–Cas9 genome editing using LNP-delivered Cas9 mRNA and sgRNA targeting the Ttr locus in hepatocytes. (b) Dose-dependent Ttr knockout efficiency following I.V. administration of Cas9 mRNA LNPs in C57BL/6 mice, showing progressive reductions in serum TTR levels. (c) Comparative evaluation of HHES-LNP and the Intellia LNP formulation LP-01, demonstrating similar levels of TTR protein reduction after systemic delivery. (d) Indel frequencies at the Ttr target site measured in liver tissue, showing comparable genome-editing efficiencies between HHES-LNP and LP-01.

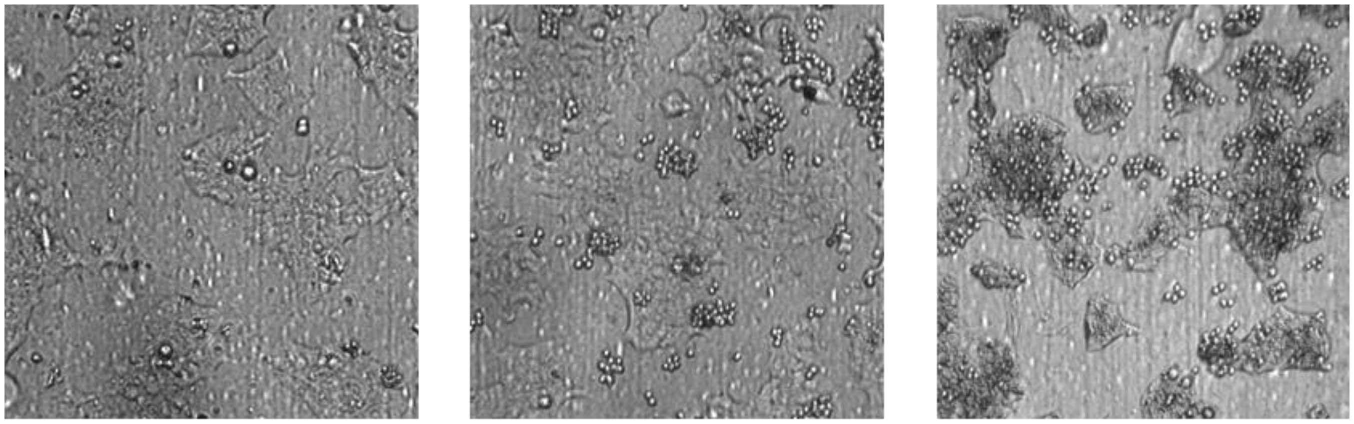


**Supplementary Figure 16. ADCC-like assay of mRNA-expressed trastuzumab.** Representative images of HER2-positive target cells following co-culture with effector cells in the presence of serum from mice treated with trastuzumab mRNA-loaded LNPs. Images illustrate target cell morphology under ADCC-like assay conditions.

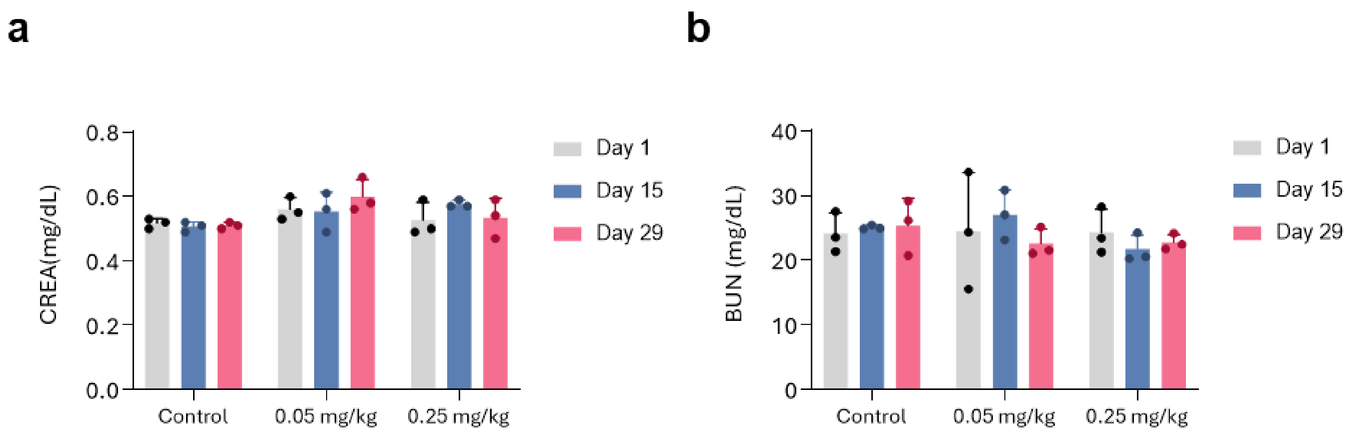


**Supplementary Figure 17. Renal function markers in non-human primates following administration of mRNA/LNPs.** Serum creatinine (CREA) (a) and blood urea nitrogen (BUN) (b) levels measured in cynomolgus monkeys after intravenous administration of mRNA-loaded LNPs. Values are presented as mean ± SEM.

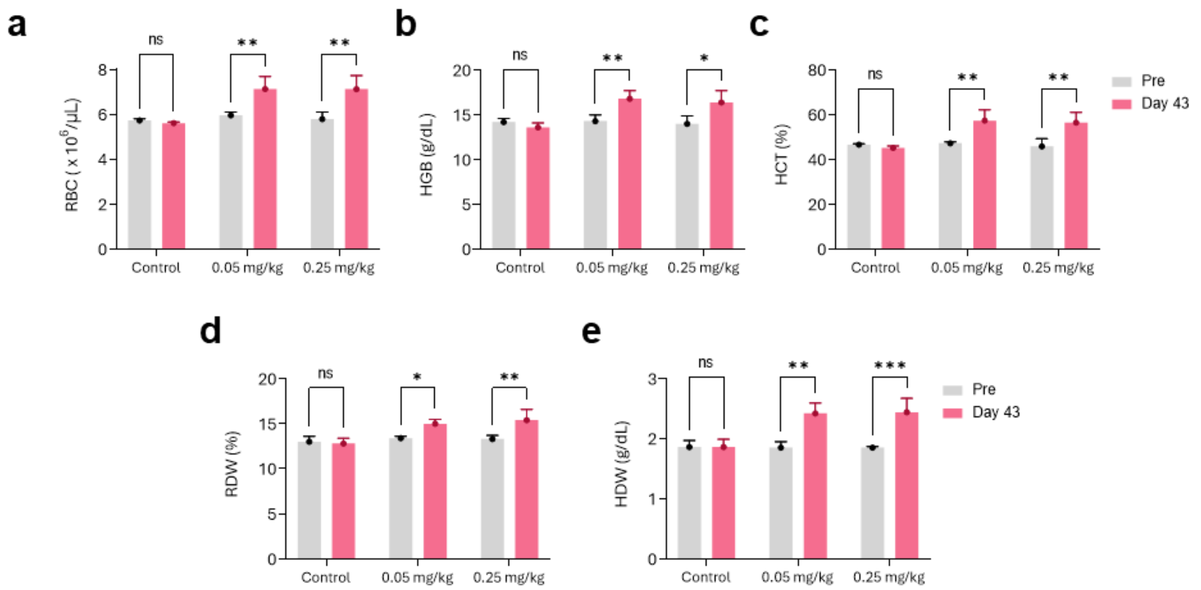


**Supplementary Figure 18. Hematological parameters following mRNA-EPO administration.** Red blood cell count (RBC), hemoglobin (HGB), hematocrit (HCT), red cell distribution width (RDW), and hemoglobin distribution width (HDW) were measured at the indicated time point. Data are presented as mean ± SD.

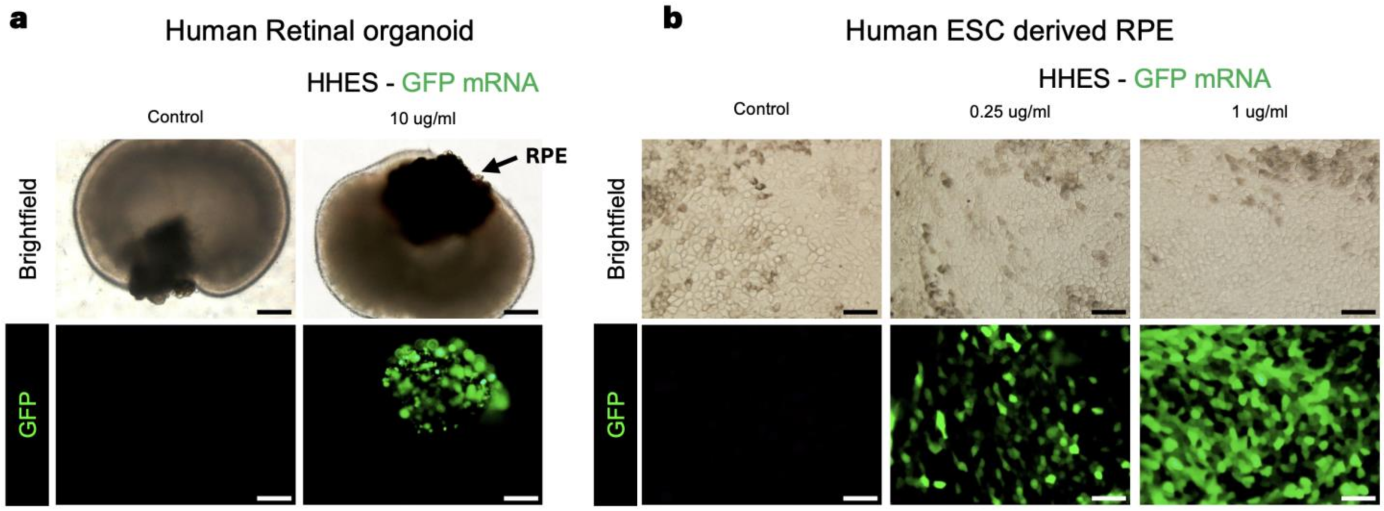


**Supplementary Figure 19. HHES LNP–mediated mRNA delivery in human retinal cell models.** (a) Representative brightfield and fluorescence images of human ESC-derived retinal organoids (>30 weeks in culture) treated with HHES-GFP mRNA LNPs (10 μg/mL) for 24 h. GFP-positive cells were detected within the pigmented RPE region, whereas no GFP expression was observed in the neural retina compartment. (b) Representative brightfield and fluorescence images of human ESC-derived RPE cells cultured as a monolayer and treated with HHES-GFP mRNA LNPs at 0.25 or 1.0 μg/mL for 24 h. Both fluorescence intensity and the proportion of GFP-positive cells increased in a dose-dependent manner. Scale bars, 200 μm in (a) and 50 μm in (b).

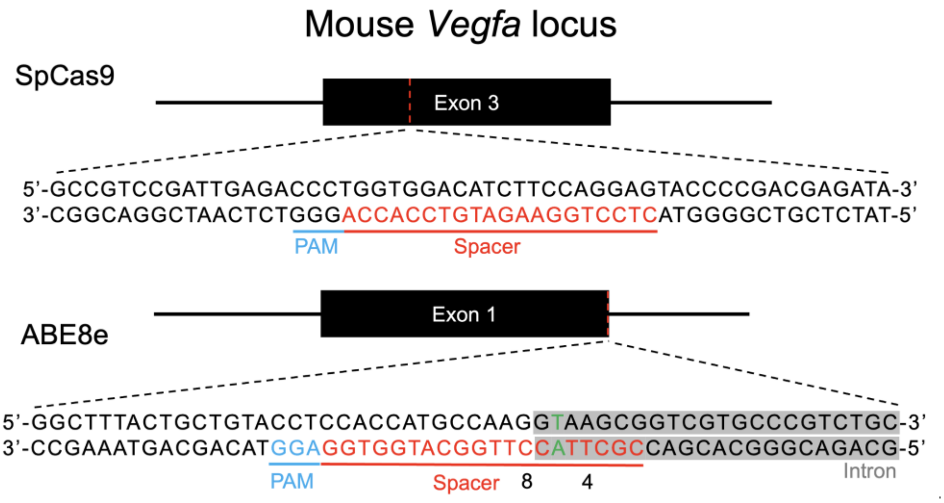


**Supplementary Figure 20. Schematic illustration of guide RNA design for Vegfa genome editing.** Diagram showing the genomic target sites used for SpCas9-mediated indel generation and ABE8e-mediated base editing at the mouse Vegfa locus. For SpCas9 editing, the sgRNA targets exon 3 (spacer: 5′-CTCCTGGAAGATGTCCACCA-3′) to induce indel formation. For ABE8e editing, the sgRNA targets the splice donor site within intron 1 (spacer: 5′-CGCTTACCTTGGCATGGTGG-3′) to disrupt canonical pre-mRNA splicing. Rosa26-targeting sgRNA (5′-GGCGGTCCTCAGAAGCCAGG-3′) served as a non-targeting control. Protospacer adjacent motif (PAM) sequences, spacer regions, and the base-editing window are indicated. The nucleotides predicted to undergo A-to-G conversion by ABE8e are highlighted in green.

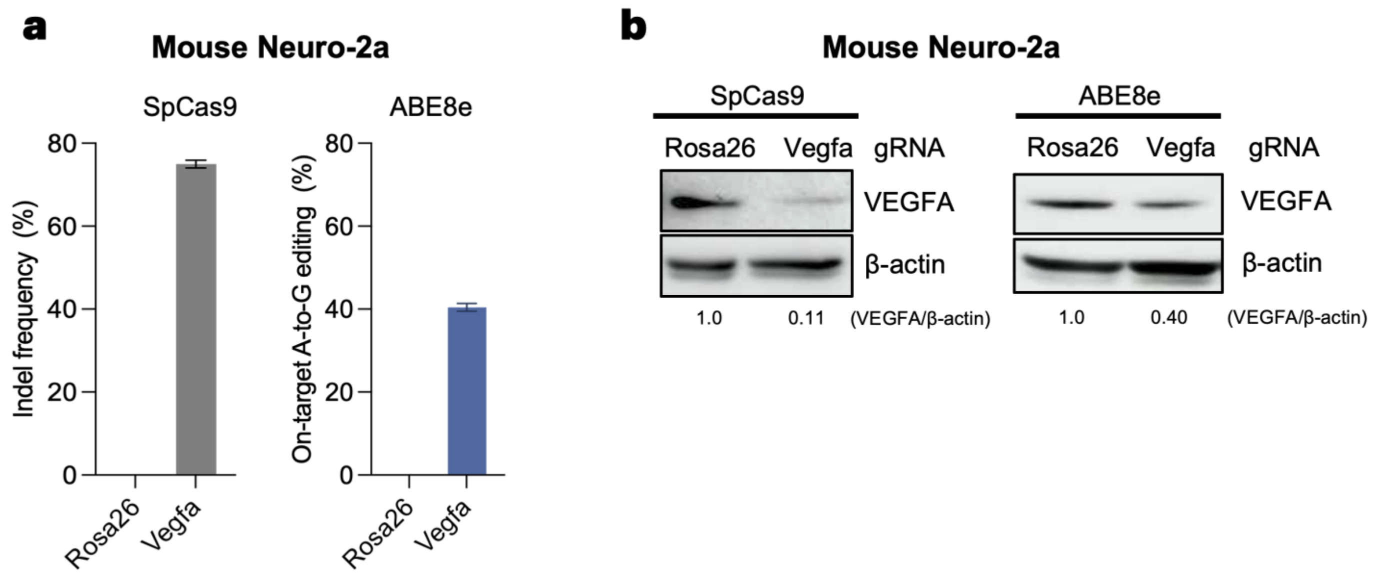


**Supplementary Figure 21. In vitro validation of Vegfa-targeting guide RNAs in mouse Neuro-2a cells.** (a) Indel frequencies for SpCas9 at the Vegfa locus and on-target A-to-G conversion efficiencies for ABE8e at the Vegfa splice donor site, quantified by targeted deep sequencing in mouse Neuro-2a cells transfected with the indicated sgRNAs. (b) Representative western blot analysis of VEGFA protein levels in mouse Neuro-2a cells transfected with SpCas9 or ABE8e together with the indicated sgRNAs. β-actin was used as a loading control. Relative VEGFA/β-actin ratios are shown below each lane, with the Rosa26 control set to 1.0.

| | Encapuslation Efficiency (%) | Size (nm) | PDI |
|---|---|---|---|
| b-2-HHES | 89.6 | 69.58 | 0.249 |
| b-3-HHES | 92.7 | 60.40 | 0.192 |
| b-4-HHES | 84.3 | 49.26 | 0.247 |

**Supplementary Table 1. Summary of physicochemical properties and formulation characteristics of LNPs.**

**a. Chemically modified sgRNA sequences**

| sgRNA | Sequence (5′-3′) |
|---|---|
| Ttr-sgRNA | 5′-mU*mU*mA*rCrArGrCrCrArCrGrUrCrUrArCrArGrCrArGrUrUrUrU<br>rArGrArGrCrUrArGrArArArUrArGrCrArArGrUrUrArArArArUrArArGrG<br>rCrUrArGrUrCrCrGrUrUrArUrCrArArCrUrUrGrArArArArArGrUrGrGrC<br>rArCrCrGrArGrUrCrGrGrUrGrCrU*mU*mU*mU-3′ |
| Vegfa-spCas9-sgRNA | 5'-mC*mU*mC*rCrUrGrGrArArGrArUrGrUrCrCrArCrCrArGrUrUrUrU<br>rArGrArGrCrUrArGrArArArUrArGrCrArArGrUrUrArArArArUrArArGrG<br>rCrUrArGrUrCrCrGrUrUrArUrCrArArCrUrUrGrArArArArArGrUrGrGrC<br>rArCrCrGrArGrUrCrGrGrUrGrCrU*mU*mU*mU-3' |
| Vegfa-ABE8e-sgRNA | 5'-mC*mG*mC*rUrUrArCrCrUrUrGrGrCrArUrGrGrUrGrGrGrUrUrUrU<br>rArGrArGrCrUrArGrArArArUrArGrCrArArGrUrUrArArArArUrArArGrG<br>rCrUrArGrUrCrCrGrUrUrArUrCrArArCrUrUrGrArArArArArGrUrGrGrC<br>rArCrCrGrArGrUrCrGrGrUrGrCrU*mU*mU*mU-3' |
| Rosa26-sgRNA | 5'-mG*mG*mC*rGrGrUrCrCrUrCrArGrArArGrCrCrArGrGrGrUrUrUrU<br>rArGrArGrCrUrArGrArArArUrArGrCrArArGrUrUrArArArArUrArArGrG<br>rCrUrArGrUrCrCrGrUrUrArUrCrArArCrUrUrGrArArArArArGrUrGrGrC<br>rArCrCrGrArGrUrCrGrGrUrGrCrU*mU*mU*mU-3' |

**Note.** In sgRNA sequences, r denotes an unmodified ribonucleotide, * denotes a phosphorothioate linkage, and m indicates a 2′-O-methyl modification.

**b. Primers for targeted deep sequencing**

| Primer | Universal 5′ adapter sequence | Locus-specific sequence (5′-3′) |
|---|---|---|
| Vegfa-spCas9-NGS-F | acactctttccctacacgacgctcttccgatct | GCCTCCGAAACCATGAACTT |
| Vegfa-spCas9-NGS-R | gtgactggagttcagacgtgtgctcttccgatct | CCCTCCACGTACGACGACAGA |
| Vegfa-ABE8e-NGS-F | acactctttccctacacgacgctcttccgatct | GGCGATAGAGGCTGACCATC |
| Vegfa-ABE8e-NGS-R | gtgactggagttcagacgtgtgctcttccgatct | CACTCCAGGGCTTCATCGTT |

**Supplementary Table 2. Chemically modified sgRNAs and primers used for targeted deep sequencing.**
